# The gut microbiota promotes liver regeneration through hepatic membrane phospholipid synthesis

**DOI:** 10.1101/2022.08.25.505228

**Authors:** Yuhan Yin, Anna Sichler, Josef Ecker, Melanie Laschinger, Gerhard Liebisch, Marcus Höring, Marijana Basic, André Bleich, Xue-Jun Zhang, Pavel Stupakov, Yasmin Gärtner, Fabian Lohöfer, Carolin Mogler, Helmut Friess, Daniel Hartmann, Bernhard Holzmann, Norbert Hüser, Klaus-Peter Janssen

## Abstract

**Background & Aims:** Hepatocyte growth and proliferation is dependent on the synthesis of membrane phospholipids. Lipid synthesis, in turn, requires short chain fatty acids (SCFA) generated by bacterial fermentation, delivered through the gut-liver axis. We therefore hypothesized that dysbiotic insults like antibiotics treatment not only negatively affect gut microbiota, but also impair hepatic lipid synthesis and liver regeneration.

**Methods:** Stable isotope labelling and 70% partial hepatectomy (PHx) was carried out in C57Bl/6J wildtype mice, in mice treated with broad-spectrum antibiotics, in germfree mice and gnotobiotic mice colonized with minimal microbiota. Microbiome was analysed by 16S rRNA gene sequencing and microbial culture. Gut content, liver and blood were tested by lipidomics mass spectrometry, qRT-PCR, immunoblot and immunohistochemistry for expression of proliferative and lipogenic markers. Matched biopsies from hyperplastic and hypoplastic liver tissue of human patients subjected to portal vein embolization were analysed by qRT-PCR for lipogenic enzymes and results were correlated with liver volumetry.

**Results:** Three days of antibiotics treatment induced persistent dysbiosis with significantly decreased beta-diversity and richness, but massive increase of *Proteobacteria*, accompanied by decreased colonic SCFA. After PHx, antibiotics-treated mice showed delayed liver regeneration, increased mortality, impaired hepatocyte proliferation and decreased hepatic phospholipid synthesis. Expression of the key lipogenic enzyme SCD1 was upregulated after PHx, but delayed by antibiotics-treatment. Germfree mice essentially recapitulated the phenotype of antibiotics-treatment. Importantly, phospholipid synthesis, hepatocyte proliferation, liver regeneration and survival were rescued in gnotobiotic mice colonized with a minimal SCFA-producing microbial community. SCD1 was required for human hepatoma cell proliferation, and its hepatic expression was associated with liver regeneration and hyperproliferation in human patients.

**Conclusion:** Gut microbiota are pivotal for hepatic membrane phospholipid synthesis and liver regeneration.

**Lay Summary:** Gut microbiota affects the liver lipid metabolism through the gut-liver axis, and microbial metabolites promote liver regeneration. Perturbations of the microbiome, e.g., by antibiotics treatment, impair the production of bacterial metabolites, which serve as building blocks for new membrane lipids in liver cells. As a consequence, hepatocyte growth and proliferation, and ultimately, liver regeneration and survival after liver surgery is impaired.

**Highlights:**

- Partial hepatectomy in mice pretreated with antibiotics is associated with impaired hepatocyte proliferation and phospholipid synthesis, delayed liver regeneration and increased mortality
- The delay in liver regeneration and impaired lipogenesis upon antibiotics treatment is preceded by dysbiosis of gut microbiota, increase of Proteobacteria and decreased short-chain fatty acids in the gut
- Partial hepatectomy in germfree mice essentially phenocopies the detrimental effects of antibiotic treatment
- Liver regeneration and mortality, as well as phospholipid synthesis and hepatocyte proliferation in germfree mice are fully rescued upon colonisation with a minimal gut bacterial consortium capable of short-chain fatty acid production
- In human patients, the intrahepatic expression of lipid synthesis enzymes positively correlates with proliferation and liver regeneration in the clinic
- Thus, liver regeneration is affected by composition of gut microbiota
- Clinically, pre-operative analysis of the gut microbiome may serve as biomarker to determine the extent of liver resections

## Introduction

The liver has a remarkable capacity to regenerate in response to partial resection or injury. Liver regeneration is a highly organized tissue regrowth process, it requires the cooperation of multiple cell types and is affected also by extrahepatic factors.[1] The crosstalk between the gut and the liver is increasingly recognized in this context,[2] since gut-derived factors like bile acids, which constantly reach the liver through the portal vein, are involved in regeneration.[3] The human gut harbors complex microbial communities, commonly referred to as microbiota, or microbiome when considering their genomes and environmental conditions. The gut microbiota has important physiological functions, such as shaping the intestinal epithelium to strengthen barrier integrity,[4] preventing the invasion of pathogens and regulating host immunity.[5, 6] Specific bacterial taxa can produce significant amounts of SCFA by fermentation from dietary fiber, which are important for immune regulation and as energy source for the host.[7–10] Moreover, we could previously show that a substantial part of lipogenesis in hepatocytes relies on microbial SCFA from the large intestine, which serve as building blocks for fatty acid synthesis.[11] Since the liver is crucial for systemic lipid metabolism, this indicates a central role of the gut-liver axis not only for liver regeneration, but also for other metabolic pathologies. To the best of our knowledge, few studies have investigated the role of gut microbiota in host lipid metabolism in the context of the regenerating liver. We hypothesized that gut microbiota provides metabolites via the portal vein that are crucial for membrane lipid synthesis, growth and proliferation of hepatocytes. Here, we investigated the putative mechanisms by which gut microbiota promote or inhibit liver regeneration. Dysbiotic insults by antibiotic treatment, inflammatory conditions of the bowel or liver cirrhosis may have profound and deleterious consequences for liver regeneration. In line, microbial signatures were reported to be associated with liver pathologies.[12–15] Specifically, gut microbial communites were significantly less diverse in patients with NAFLD-cirrhosis, Firmicutes were decreased, while Proteobacteria were enriched. Of note, liver disease is still a major cause of worldwide premature mortality in adults.[16]

Further, hepatectomy is an effective oncologic treatment for primary and metastatic liver malignancies and can provide the best survival outcome in many clinical situations. However, many patients require extensive hepatic resection due to the large size of the tumour(s) adjacent to important vascular structures, resulting in an insufficient volume of the remaining liver after radical resection. Thus, ensuring adequate future liver remnant (FLR) volume and functionality is a key factor in hepatic tumour resection. To address how gut-derived bacterial metabolites contribute to different phases and aspects of liver regeneration, we have used partial hepatectomy (PHx) as preclinical surgical model.[17] We found that treatment with broad-spectrum antibiotics for three days prior to PHx caused severe microbial imbalance, accompanied by significant decrease in short-chain fatty acids (SCFA) in the gut. Hepatocytes in antibiotics-treated mice showed strongly decreased proliferation and impaired phospholipid synthesis, accompanied by reduced hepatic expression of the enzyme SCD1, which catalyzes the rate-limiting step in the formation of monounsaturated fatty acids.[18] Liver regeneration was significantly impaired and delayed upon antibiotics treatment and associated with increased mortality. In order to tackle the functional importance of bacterial-derived SCFA for hepatic lipid synthesis, proliferation and regeneration, gnotobiotic as well as germfree mice were treated with PHx. Compared to complex microbiota colonized controls (i.e., “conventionally” housed), germfree mice closely reproduced the phenotype observed upon antibiotics treatment. Liver regeneration and mortality were rescued by colonizing gnotobiotic mice with a minimal microbial community capable of SCFA-production. SCD1 was found to be required for cell proliferation in human hepatoma cells. Furthermore, expression of SCD1 was tested in matched patient tissue biopsies from hyperplastic and hypoplastic liver tissues after surgical intervention to induce liver growth, and a positive correlation with liver regeneration was found. Taken together, gut microbiota play a crucial role in lipid metabolism to promote liver regeneration, and gut dysbiosis can lead to impaired liver regeneration.

## Materials and Methods

### Mice

All animals received human care, all mouse experiments were conducted in accordance with the EU directive 2010/63 and the German animal welfare law and all other relevant national and institutional guidelines. Female C67BL/6J mice aged 10- 12 weeks in several groups were analysed: specific-pathogen-free (SPF), germfree (GF) and Oligo-Mouse-Microbiota12 (OMM),[19]. Mice were obtained from the Institute for Laboratory Animal Science (Hannover Medical School, Hannover, Germany). SPF mice were housed in individually ventilated cages, OMM and GF mice were housed in sterile static microisolators (Han-gnotocages) with HEPA-filters to ensure gnotobiotic housing conditions. All mice were maintained in a controlled environment with a 12h light–dark cycle and received sterile 50 kGy gamma-irradiated standard diet (V1124-927, Ssniff, Soest, Germany), and autoclaved water ad libitum. Sterility of GF mice was routinely confirmed by culturing and microscopic observation of feces after Gram-staining. In addition, 16S rRNA gene-targeted PCR of cecal content from all mice was performed at the end of the study. The experimental use of mice in was approved by local authorities (ROB license 55.2-1- 54-2532-147-2015).

### Antibiotics Treatment

Studies were performed in female C57BL/6 mice, aged 10-12 weeks, under SPF conditions housed in a barrier facility (ZPF, Klinikum rechts der Isar, TUM), in individually ventilated cages in a controlled environment with 12 h light–dark cycle and received ad libitum autoclaved standard feed (no. 1324SP, Altromin, Lage, Germany). For antibiotics treatment, mice received 1 g/L Ampicillin, 0.5 g/L Vancomycin, 1 g/L Metronidazol, 0,2 mg/L Fluconazol (all from Sigma-Aldrich, Taufkirchen, Germany), and Splenda sweetener at 1.2% (weight/volume; TC Heartland, LLC, USA) according to published protocols[20], in autoclaved drinking water for 3 days before PHx and continued until sacrificed. Control mice received autoclaved drinking water with 1.2% Splenda for the same duration. Drinking water for both groups were changed every three days.

### Partial hepatectomy

Partial hepatectomy (PHx) was performed in mice from 10 a.m. to 12 a.m., as described.[17] Briefly, mice were anesthetized with isoflurane, ligation and resection of the median lobe and the left lateral lobe was performed separately. GF and OMM mice were operated under sterile laminar flow, and placed back in sterile microisolators after the intervention. Animals were sacrificed after indicated time periods (Fig. 1A, ≥6 mice per group and timepoint), or when humane endpoints were reached. Liver, blood, colon content and feces of mice were processed immediately or shock frozen in liquid nitrogen.

**Fig. 1.**
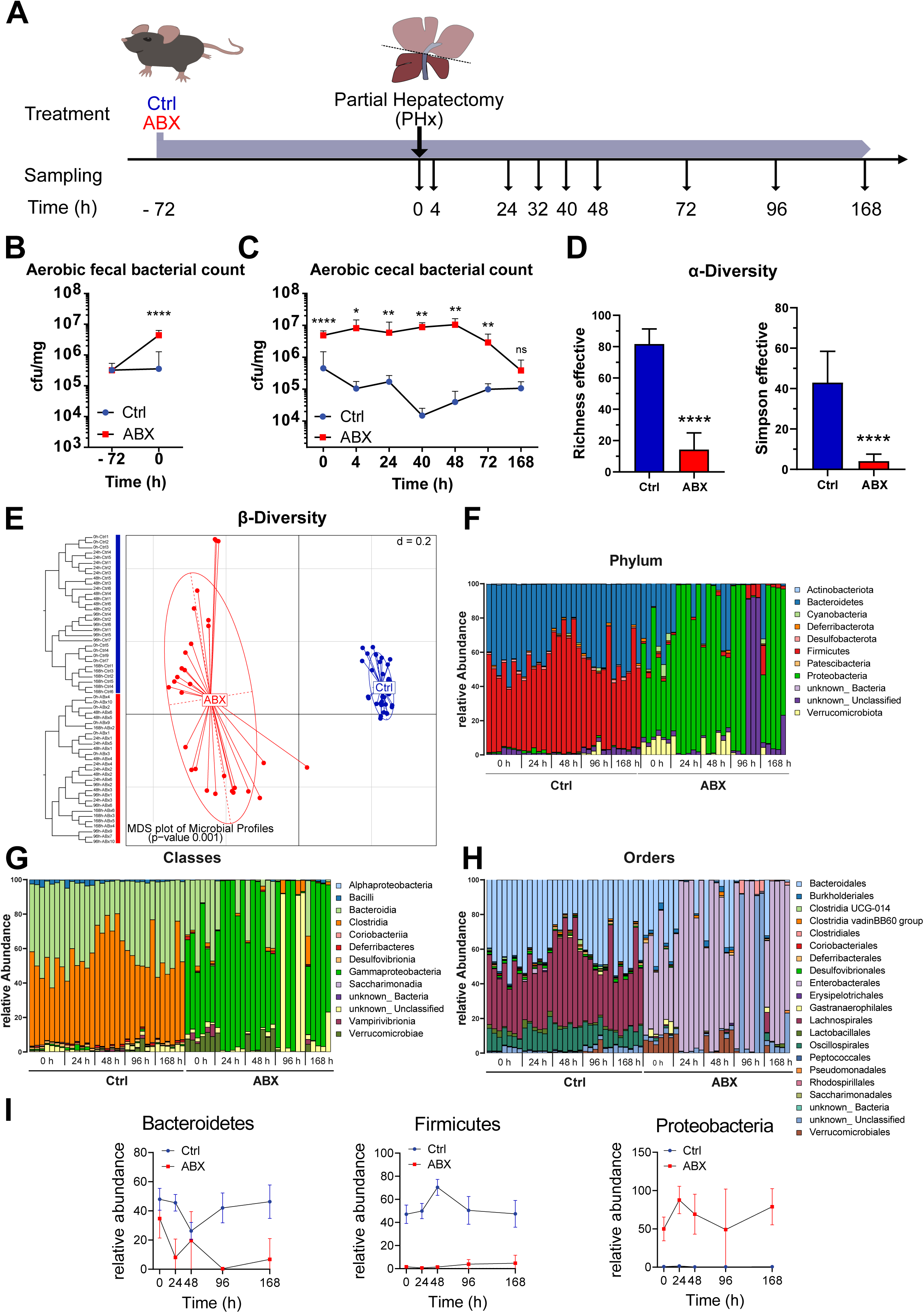
Treatment with broad spectrum antibiotics induces gut dysbiosis. (A) Treatment with broad spectrum antibiotics (ABX, n=68) or control (Ctrl, n=59) drinking water started three days prior to partial (2/3) liver resection (PHx), and continued until the endpoint. Sampling timepoints are indicated with small arrows. (B) Number of aerobically grown colony forming units (cfu) per mg of feces before and after ABX or Ctrl treatment. (C) Aerobically growing cfu/mg of cecal content at sampling. Displayed is mean ±SD, significance was determined by Mann-Whitney U test where P>0.05 = ns, P≤0.05 *; P≤0.01 **; P≤0.001 ***; and P≤0.0001 ****. (D) Diversity of microbial communities within individual samples (α-diversity) based on V3-4 16S rRNA sequencing at all timepoints, displayed as mean effective richness and Simpson effective coefficients (±SD), unpaired two-tailed student’s-t test. (E) β- Diversity differs significantly between ABX and control groups in multidimensional scaling (MDS) plot. Relative abundances of taxa at phylum (F), class (G) and order level (H) are drastically altered between ABX and Ctrl. (I) Relative abundance of the main phyla Bacteroidetes, Firmicutes, and Proteobacteria over time.

### Liver histology and immunohistochemistry

Liver samples were fixed in 4% paraformaldehyde for 72 hours followed by dehydration and embedding in paraffin. Rehydrated tissues sections of 3 µm were incubated with anti-Ki67 antibodies (BD Pharmingen) and staining was visualized using Dako EnVision+ (Agilent Technologies, Santa Clara, CA). Positively stained hepatocyte nuclei in five random high-power microscopic fields were counted for quantification. Hematoxylin & eosin (H&E) staining was conducted according to standard protocols.

### Immunoblotting

Protein from shock-frozen liver samples was isolated in lysis buffer containing 1% Triton X100, 150 mM NaCl, 20 mM Tris-HCl, pH 7.5, protease and phosphatase inhibitors (PMSF, Aprotinin, Pepstatin, Leupeptin) and homogenised with autoclaved metal beads in a TissueLyser II instrument (Qiagen, Hilden, Germany), followed by ultrasonication (8 sec, repeated four times) on ice, and centrifugation for 15 min at 13.000 rpm 4°C. 10 µL lysate per lane (corresponding to 20-40μg protein) were separated by denaturing SDS-PAGE and transferred to a nitrocellulose membrane. Membranes were blocked and incubated with the primary antibodies overnight, washed and incubated with secondary antibody goat-anti-rabbit-HRP or goat-anti-mouse-HRP (Promega) for 1 h. Antibodies are shown in detail in supplementary table 1. Signals were visualized using Pierce ECL western blotting detection system (Thermo Fisher Scientific) and a ChemStudio Plus instrument (Analytik Jena, Jena). Densitometric analyses and qualification were performed using ImageJ software (https://imagej.nih.gov/ij/).

### Quantitative reverse transcriptase PCR

Total RNA was extracted from snap frozen mouse or human liver tissue using NucleoSpin RNA Mini kit (Macherey-Nagel, Düren, Germany). First strand cDNA was synthesized from 1 μg total RNA using the QuantiTect Reverse Transcription Kit (Qiagen). RT-qPCR was performed in a LightCycler480 (Roche, Mannheim, Germany) using KAPA SYBR FAST Kit (KK4611, Sigma-Aldrich) or Universal Probe Library (Roche). Oligonucleotide primer sequences are shown in detail in the supplementary table 2. Relative mRNA expression was quantified normalizing against housekeeping transcripts GAPDH or ACTB (β-actin). For human transcripts, RNA was isolated following previously described protocol.[21]

### Cell culture and proliferation assay

Human cell line HepG2 was purchased from American Tissue Culture Collection (ATCC, Manassas, VA) and cultured in Dulbeccòs Modified Eagle Medium (Thermo Fisher Scientific), 1% L-glutamine with 10% fetal bovine serum, and 1% penicillin/streptomycin in 5% CO2 at 37 °C. The cells were transfected with SCD siRNA (sc36464, Santa Cruz, CA, USA) using Lipofectamine RNAiMAX (Invitrogen, Carlsbad, CA, USA) according to the manufacturer’s instructions. After 48 hours, cells were harvested for further analysis. Proliferation was determined using an MTT assay (#M5655, Sigma-Aldrich). Cells were seeded at a density of 2×10^3^ cells per well in 96-well plates with complete medium 24 h prior to treatment with 10 µM of a selective SCD1 inhibitor dissolved in DMSO (ab142089, Abcam) or an equal amount of DMSO vehicle for various time points. Then, MTT reagent was added and absorbance measured at 560 nm in a Microplate reader (Berthold, Bad Wildbad, Germany), according to the manufacturer’s instructions.

### Statistical analysis

All data are expressed as mean ± SD unless indicated differently. Statistical differences were analysed using two-tailed unpaired Student’s t test or Mann-Whitney test. Differences with p ≤ 0.05 were considered statistically significant with P>0.05 = ns, P≤0.05 *; P≤0.01 **; P≤0.001 ***; and P≤0.0001 ****. All correlations were calculated in GraphPad Prism 8.0 for Spearman’s or Pearson’s correlation coefficient, respectively. Pearson’s correlation is a measure of a linear correlation in the data, while Spearman’s correlation coefficient is based on ranked values rather than the values themselves. For all mouse experiments animal sample sizes were calculated in advance using power analysis and based on personal experience; the group allocation of the animals was not blinded (practically impossible), but mice were randomly chosen for the different antibiotic treatments.

### Clinical specimens

Three patients from the Dept. of Surgery, TUM, with liver metastasis from bile duct cancer, colon carcinoma or squameous cell carcinoma of the bile duct were included. All experiments were approved by the ethics committee of the Faculty of Medicine of TUM (vote 86/17S). Written informed consent of the patients for the use of tissues was obtained according to the ethical guidelines of the World Medical Association (WMA) Declaration of Helsinki. Liver tissue samples were obtained from each patient within (a) the hyperplastic and (b) the hypoplastic liver lobe post-surgical intervention to induce hyperplasia. In order to calculate the regeneration index within each lobe liver segmentation was performed using the segmentation software (IntelliSpace Portal Version 12.1; Phillips Medical Systems, Eindhoven, The Netherlands) on diagnostic CT scans. Functional liver volume was segmented at three consecutive time points: prior to surgical intervention, after intervention, and after resection. Tumour volume was excluded from the segmentation.

### In vivo stable isotope labelling, lipid species quantification and metabolic profiling

Lipids were extracted according to the procedure described by Bligh and Dyer in the presence of non-naturally occurring lipid species as internal standards.[22] Quantification was performed by flow injection analysis electrospray ionization tandem mass spectrometry (ESI-MS/MS) in positive ion mode using the analytical setup and strategy as described.[23] A fragment ion of m/z 184 was used for phosphatidylcholine (PC), sphingomyelin (SM) and lysophosphatidylcholine (LPC).[23] The following neutral losses were applied: Phosphatidylethanolamine (PE) 141, phosphatidylserine (PS) 185, phosphatidylglycerol (PG) 189 and phosphatidylinositol (PI) 277, according to published protocols [24]. PE-based plasmalogens (PE P) were analyzed according to principles described.[25] Sphingosine based ceramides (Cer) and hexosylceramides (HexCer) were analyzed using a fragment ion of m/z 264.[26] At 2h before the respective sampling time point, [D9]-Choline, or [D9]-Choline and [D4]-Ethanolamine (both Sigma-Aldrich), dissolved in 0.2 ml sterile NaCl solution (0.9%) was applied via i.p. injection. [D9]- Choline labelled lipids were analyzed by a fragment ion of m/z 193, PE[D4], and PC[D4] by a neutral loss of 144 and fragment ion of m/z 188, respectively.[27] Self-programmed Excel Macros performed correction of isotopic overlap of lipid species as well as quantification using calibration lines as described previously.[23, 28] Lipid species were annotated according to the recently published update for shorthand notation of lipid structures.[29]. Glycerophospholipid species annotation was based on the assumption of even numbered carbon chains only.

For analysis of short chain fatty acids, frozen colon content samples were homogenized in 70%-isopropanol by bead beating (Precellys, Bertin, Montigny-le-Bretonneux, France). Short chain fatty acids were then quantified by LC-MS/MS as described in detail.[30]

### 16S rRNA gene amplicon sequencing and analysis

Cecal content was prepared and analysed as previously described.[31] Briefly, cecum content was snap frozen, DNA stabilizer was added, and isolation was performed by mechanical disruption and chemically followed by NucleoSpin gDNA Clean-up Kit (Macherey-Nagel). RNA was removed with ribonucleases and genomic DNA concentration was quantified with NanoDrop. Hence the amplicon library was constructed on basis of V3-V4 region of genomic16S rRNA PCR, followed by amplicon cleaning, dilution and sequencing with Illumina MiSeq as published.[32] Data analysis was performed with the platforms Integrated Microbial Next Generation Sequencing (IMGNS) and Rhea.[33]

### Aerobic cultivation of gut bacteria

Bacteria in mouse caecum or feces were cultivated on blood agar plates (Inst. of Med. Microbiology, Immunology and Hygiene, TUM) under sterile conditions. Fresh samples were weighed and transferred into 2 ml Eppendorf tubes containing 1 ml sterile PBS and an autoclaved metal bead, followed by vortexing for 15 sec. Serial dilution was performed until a 10^-7^ dilution was reached. Sample homogenization was ensured by vortexing for 3 sec before the respective volume was taken out. Four serial dilutions were plated, and bacteria were cultivated overnight at standard aerobic conditions at 37°C. Single colonies were counted and colony forming units (cfu) per mg were calculated.

## Results

### Antibiotics treatment during partial hepatectomy induces dysbiosis

The role of the microbial composition in the course of liver regeneration, as well as the consequences of a dysbiotic insult were studied in a mouse model of 70% partial hepatectomy (PHx). Dysbiosis was induced by three days of broad-spectrum antibiotics administered in drinking water prior to PHx. Treatment with antibiotics (ABX, n=68 mice) or water supplemented with the sweetener splenda (control, n=59 mice) was continued until animals were sampled at indicated timepoints (Fig. 1A). Efficacy of treatment was confirmed by aerobic culture of fecal pellets and cecal content, and by 16S rRNA sequencing of cecal content. Antibiotics had no detrimental effects on general health or body weight (not shown), but resulted in a significantly increased number of aerobic colony forming units in feces collected from live mice prior to PHx (Fig. 1B). In accordance, a significant increase in aerobic colony-forming units was observed in cecal contents in the antibiotics-treated group compared to controls, observable until 72 h after PHx (Fig. 1C). As expected, 16S rRNA amplicon sequencing indicated a highly significantly decreased individual microbial diversity (α-diversity), depicted as richness-effective and Simpson-effective coefficients (Fig. 1D). The β- diversity differed significantly between both groups (Fig. 1E), and the heterogeneity within the antibiotics-treated group was far greater than in controls. Taxonomic analysis showed a shift from Firmicutes and Bacteroidetes to a Proteobacteria-dominated community under antibiotics-treatment (Fig. 1F). On taxonomic class level, Clostridia were almost absent in antibiotics-treated mice, Bacteroidia were reduced, whereas Gammaproteobacteria showed the highest abundance (Fig. 1G). Regarding taxonomic order, Lachnospirales and Oscillospirales were absent and Bacteroidales reduced in antibiotics-treated mice (Fig. 1H). Relative abundance of Bacteroidetes remained relatively stable in controls (Fig. 1I), but decreased over time in antibiotics-treated mice. Firmicutes are highly present in control mice with few changes over time, but almost absent in the antibiotics-treated group (Fig. 1I). In contrast, the relative abundance of Proteobacteria was very low in controls, but strongly increased in antibiotics-treated mice. Moreover, there are strong alterations between the samples in the antibiotics-treated group, indicating unstable microbiomes (Fig. 1I).

### Antibiotics treatment is associated with decreased liver regeneration, survival and hepatocyte proliferation

To evaluate whether liver regeneration is affected by a dysbiotic insult due to broad spectrum antibiotics treatment, liver to body weight ratio of mice was determined after PHx, and a highly significantly decreased liver to body weight ratio was detected at most time points (Fig. 2A). A slight but significant reduction in liver weight ratio was found in antibiotics treated mice prior to PHx (Fig. 1 and Suppl. Fig. 1A). After normalizing to the initial liver weight at the time of hepatectomy in controls and antibiotics-treated mice, respectively, liver/body weight ratios were still substantially reduced upon antibiotics treatment after PHx compared to controls (Suppl. Fig. 1A, B). Moreover, antibiotics-treated mice showed significantly higher mortality, defined as reaching the humane endpoint (Fig. 2B). Antibiotics-treated mice displayed a significant delay in the number of proliferating hepatocytes, evidenced by the marker Ki67 that was significantly decreased between 32h and 72h after PHx (Fig. 2C, D, Suppl. Fig. 1C). Next, critical cell cycle regulators were assessed. In accordance with Ki67 stainings, mRNA levels of cyclin D1 (Fig. 3A) and cyclin E1 (Fig. 3B), associated with late G1-phase, were found to be delayed in antibiotics-treated mice. Cyclin A2 (Fig. 3C) and cyclin B1 (Fig. 3D), effective in S- and G2-phase, respectively, showed delayed activation in antibiotics-treated mice. Interestingly, cyclins E1 and A2 showed a compensatory increase of expression in antibiotics-treated mice at the 168h timepoint. On protein level, phosphorylation of retinoblastoma protein (p-RB), required for transition beyond the G1/S restriction point, total expression of cyclins A2, B1, and the cell-cycle dependent kinase CDK1 were tested (Fig. 3E). Immunoblot analysis confirmed a generally decreased expression of cell cycle regulators until 48-72h post PHX in antibiotics-treated mice. In accordance with the mRNA expression, there was an increase of cell cycle regulators visible at the late timepoint of 168h in antibiotics-treated mice, as shown by quantification of immunoblots (Fig. 3F-I). To assess cell growth (i.e., volume increase) in addition to cell proliferation, hepatocyte cell sizes were quantified histologically. Antibiotics-treated mice did not show an increase in hepatocyte cell volumes compared to controls; actually, apparent hepatocyte size was reduced in antibiotics-treated mice, differences attaining significance at 32h and 48h timepoints (Suppl. Fig. 1D).

**Fig. 2.**
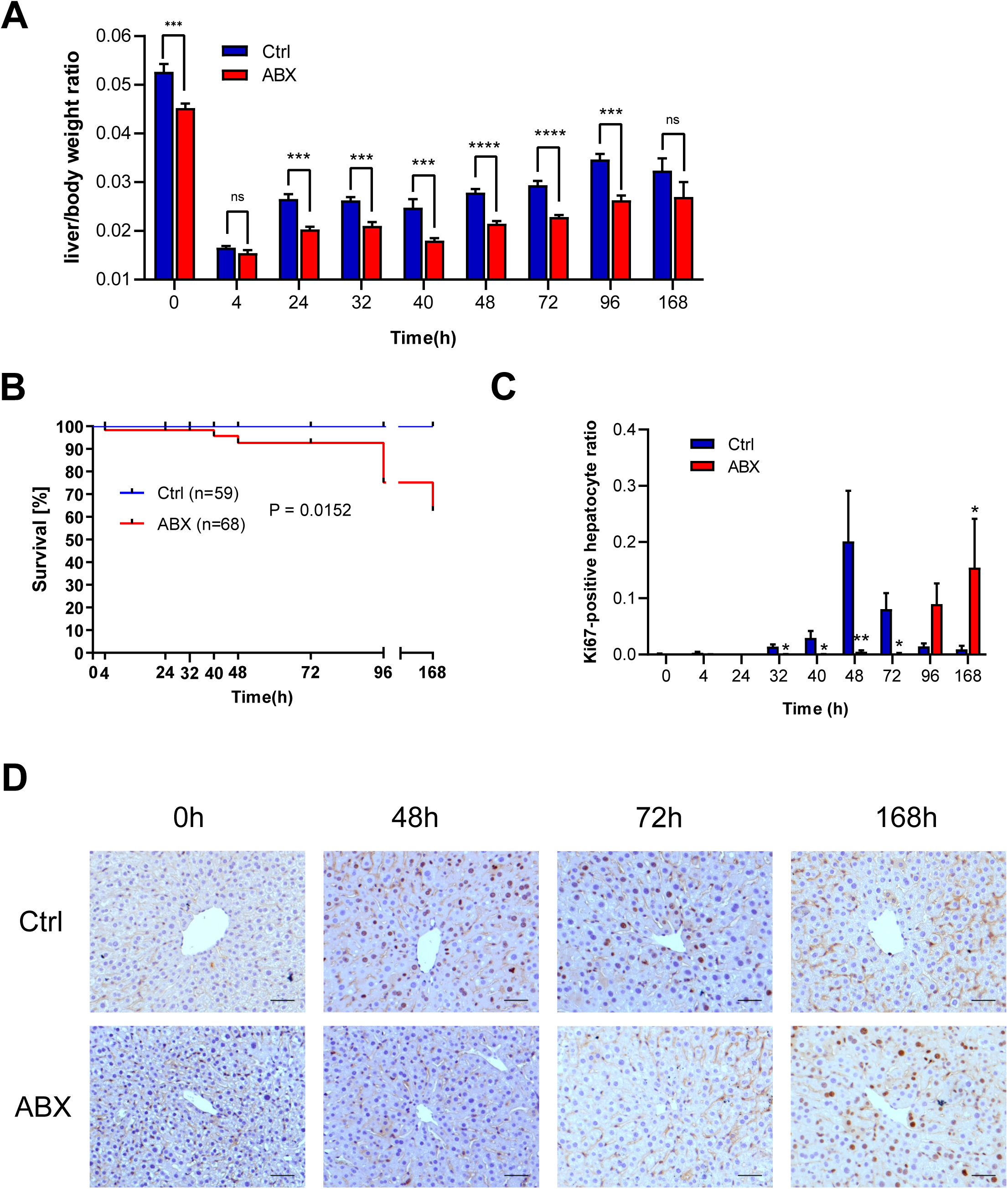
Antibiotics-treated mice show impaired liver regeneration, survival and hepatocyte proliferation after PHx. (A) Liver/body weight ratio was determined at indicated timepoints (n=6-10 mice per timepoint and group). (B) Kaplan-Meier analysis of the survival rate in antibiotics- treated mice and controls after hepatectomy. (D-E) Quantification of Ki67-positive hepatocytes and representative immunohistochemical images are shown. Scale bars represent 50 µm. Data were analysed using the two-tailed Student’s t-test or the Mann- Whitney U test. Data are presented as mean ± SEM.

**Fig. 3.**
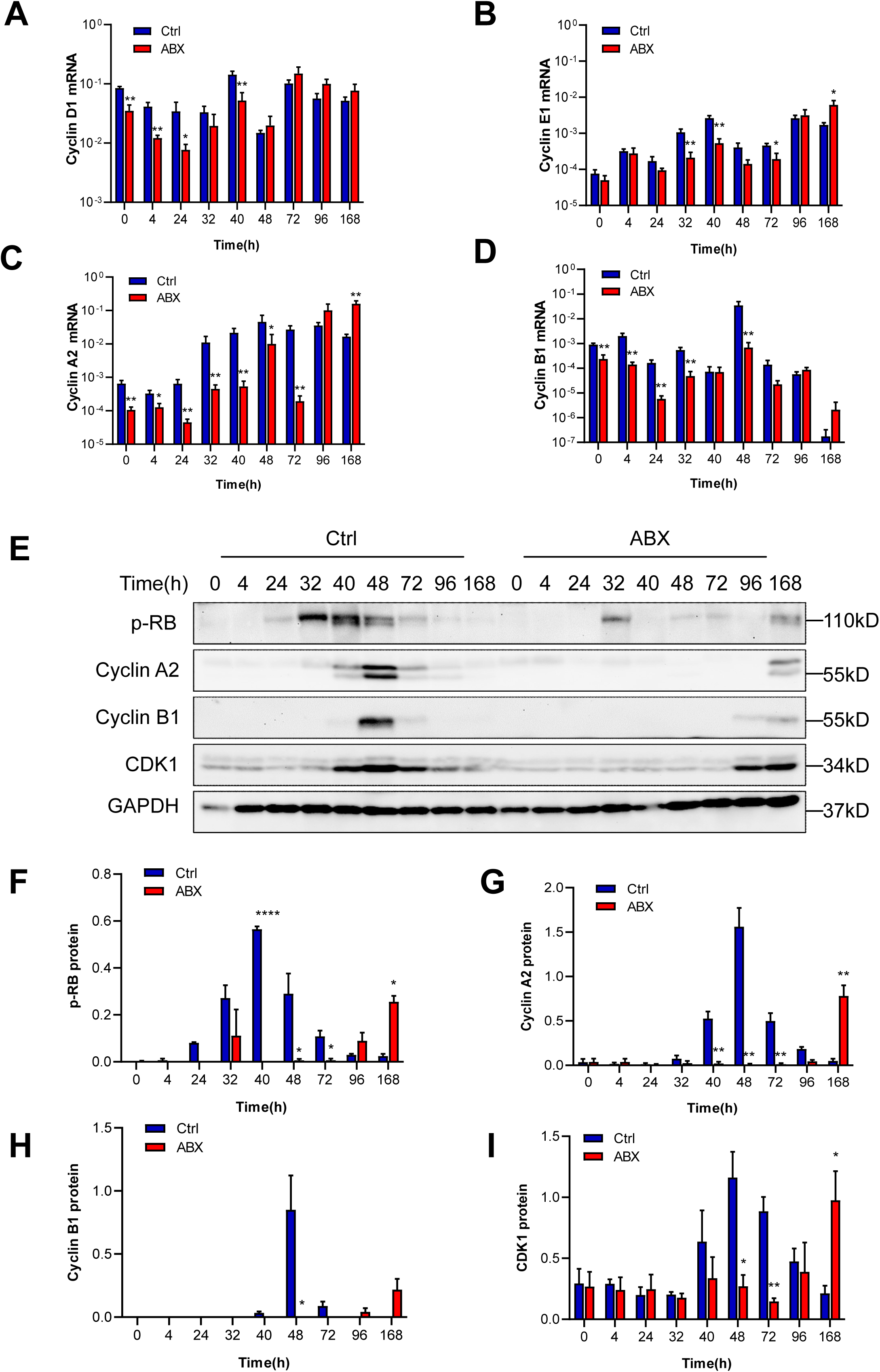
Hepatic cell cycle progression after PHx is generally delayed in antibiotics- treated mice. (A-D) Hepatic mRNA expression of cyclins D1, E1, A2 and B1 were normalized to those of housekeeping transcript GAPDH and compared between antibiotics-treated (ABX) and control mice. (E - I) Protein expression of phosphorylated RB, Cyclin A2, Cyclin B1 and CDK1 in mouse liver after PHx were analysed by immunoblot. Representative blots (E) and densitometric analyses (F, G, H and I) are depicted. Data from three or more independent experiments were analysed using two-tailed student’s t test or Mann-Whitney U test. Data are presented as mean±SEM.

### SCFA depletion in gut content due to antibiotic treatment

As depicted schematically (Fig. 4A), the influence of gut microbiota on hepatic lipid metabolism was assessed on several levels, by labelling with the stable isotope phospholipid precursor [D9]-choline, administered by intraperitoneal injection.

**Fig. 4.**
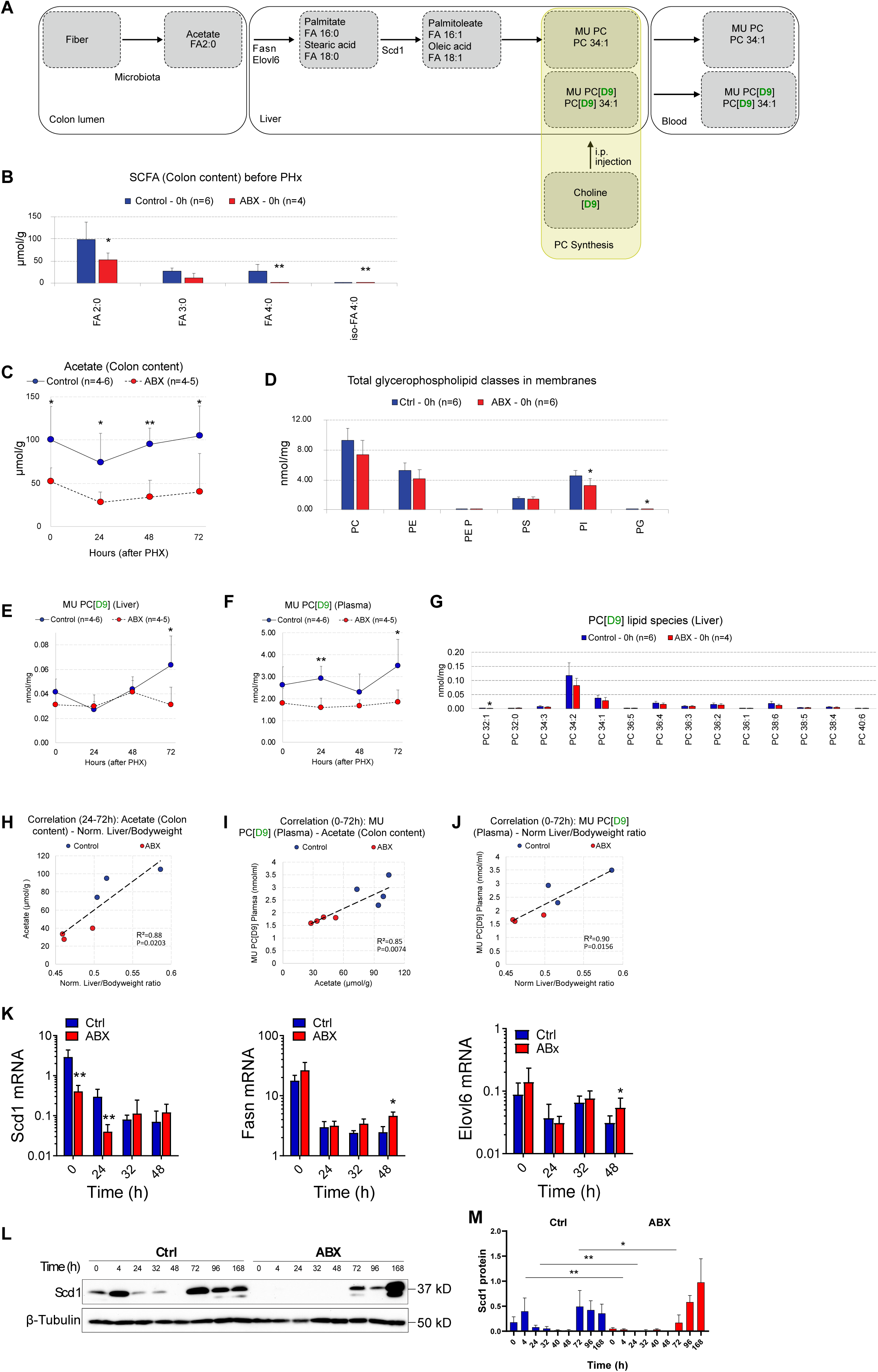
Short chain fatty acid content in gut and hepatic lipid synthesis are decreased in antibiotics-treated mice. (A) Schematic depiction of fiber being transformed by bacterial fermentation in the gut lumen to short-chain-fatty-acids (SCFA), e.g., acetate (FA 2:0), which reach the liver via the portal vein and are there converted to saturated fatty acids (FA). These FA are desaturated by SCD1 to monounsaturated fatty acids (MUFA), and incorporated in phospholipids (e.g., in phosphatidylcholine as MU-PC species 34:1). De novo lipid synthesis is monitored through i.p. injection of stable isotope labeled D9-choline, yielding monounsaturated phosphatidyl choline (MU-PC[D9]), which can be measured in liver or blood. (B) Quantity of short chain fatty acids (SCFA) acetate (FA 2:0), propionate (FA 3:0), butyrate (FA 4:0), and iso-butyrate (iso-FA 4:0) in colon content of mice in ABX (red) compared to control mice (blue). (C) Acetate in colon content over time. (D) Quantity of glycerophospholipids (GPL), with phosphatidylcholine (PC) as the main lipid class, is essentially conserved in ABX mice. (E - F) Newly synthesized MU PC[D9] in liver (E) or blood plasma (F). (G) Newly synthesized lipid species after three days ABX treatment displayed as PC coupled to fatty acids (FA) with the respective number of C-atoms measured by GC-MS/MS and indicated number of double bonds. (H) Correlation of normalized liver/bodyweight ratio and acetate in colon content at 24h, 48h and 72h (I) Correlation of de-novo synthesized MU PC[D9] measured in blood and acetate in colon content (I) or normalized liver/bodyweight ratio (J) at 0h, 24h, 48h and 72h after PHx. (K) Hepatic mRNA expression of lipogenic enzymes SCD1, FASN and ELOVL6. (L) Kinetics of Scd1 expression on protein levels, normalized to β-tubulin, and respective quantification (M). Displayed is mean±SEM, significance was tested using Mann-Whitney U test with significances as indicated above.

Briefly, important groups of microbiota, such as Firmicutes and Bacteroidota, but not Proteobacteria, are well known to produce SCFA, mainly acetate (FA 2:0), from dietary fiber by fermentation in the gut. SCFA reach the liver via the portal vein and are incorporated there in de novo synthesized saturated fatty acids (FA) through host enzymes FASN (fatty acid synthase). The resulting FA are elongated by the enzyme ELOVL6, and/or processed by the desaturase SCD to yield monounsaturated fatty acids (MUFA). MUFA are incorporated into phospholipids, such as phosphatidylcholine (PC), as most abundant phospholipid class in mammalian cell membranes, with PC 34:2 as the most abundant lipid species. Lipogenesis in the liver is crucial for systemic metabolism, as evidenced by analysis of blood serum samples. We found all SCFA to be significantly reduced in the gut of antibiotics-treated mice prior to hepatectomy (Fig. 4B, Suppl. Fig. 2A-D). Among the SCFA, acetate is the most abundant species (FA 2:0), it serves as central building block for FA synthesis in the liver. Of note, during liver regeneration, acetate levels in the colon remain significantly lower in antibiotics-treated mice (Fig. 4C). As expected, global composition of glycerophospholipids in the liver does not differ in a major way between antibiotics-treated mice and controls. Interestingly, however, phosphatidyinositol (PI) and phosphatidylglycerol (PG) were significantly reduced in antibiotics-treated mice. Next, we used stable isotope labeling to directly monitor de novo hepatic synthesis of membrane lipids. In control mice, the sum of [D9]-labelled PC (carrying nine deuterium atoms) in the liver showed a clear increase after PHx, indicative of a demand for newly synthesized lipids in proliferating hepatocytes (Fig. 4E). This increase was blunted in antibiotics treated mice, attaining significance at 72h timepoint. Accordingly, [D9]-labelled PC levels were significantly lower in blood plasma (Fig. 4E, F) in antibiotics-treated mice. This indicates that the increase in lipid synthesis in the liver during liver regeneration is absent in antibiotics treated mice. Next, the distribution of PC species was analysed in liver samples, and a trend towards more saturated species was notable in antibiotics treated mice, with similar overall species profiles (Fig. 4G). Next, we tested whether the observations in gut and liver were actually associated, in order to test the initial hypothesis SCFA produced by gut bacteria are required for phospholipid synthesis in the liver, and, ultimately, liver regeneration. The concentration of acetate, the most abundant SCFA in gut content, was significantly and positively correlated to the liver/body weight ratio (Fig. 4H). Further, newly synthesized PC, measured in blood plasma, was significantly correlated with acetate concentrations in the colon (Fig. 4I), as well as with liver/body weight ratios (Fig. 4J). Prominent monounsaturated PC species (32:1, 34:1, 36:1) were significantly decreased in liver tissue (Suppl. Fig. 2G-I), as well as in blood plasma (Suppl. Fig. 2J-L). Moreover, correlation analyses showed a positive correlation between gut SCFA concentrations, hepatic and systemic PC levels, liver-to-body-weight ratio and cyclin E1 (Suppl. Fig. 2E, F, M-R).

We further assessed expression of key enzymes involved in fatty acid synthesis and modification. The desaturase SCD1 was decreased upon antibiotics-treatment during early phases of regeneration, both on mRNA (Fig. 4K) and protein level (Fig. 4L). The fatty acid synthase FASN and the elongase ELOVL6, in contrast, showed a stronger upregulation in antibiotics treated mice (Fig. 4K). Further, SCD1 showed a clear upregulation on protein levels at two phases of liver regeneration, at very early (4h) and later (72h) timepoints. The first peak was completely absent in antibiotics treated mice, and SCD1 expression delayed until 168h (Fig. 4M). Of note, expression kinetics of SCD1 in control and antibiotics-treated mice was reminiscent of the cell cycle regulators, which were delayed to the 168h timepoint in antibiotics-treated mice (Fig. 3E).

### Germ-free mice fail to regenerate after partial hepatectomy, colonization with a minimal bacterial consortium rescues liver regeneration

In order to test the impact of fermentation-competent microbiota on liver regeneration, PHx was carried out in a complementary and independent model. Complex microbiota colonized mice (SPF, specific pathogen free, n=7) were compared to germfree mice (GF, n=5). Importantly, both groups were of the same age, sex and genetic background, bred and maintained in the same facility, and received the same sterilized, germfree food. Mice were sacrificed and sampled at 48 h after PHx (Fig. 5A). In GF mice no microbiota was detectable prior to PHx and at 48h, based on lack of 16S rRNA gene amplification (Suppl. Fig. 3A). Consequently, species diversity as well as taxonomical abundance measures cannot be displayed (n.d., not detectable) (Fig. 5B - D). A drastically higher number of GF mice reached the humane endpoint after PHx compared to SPF mice. This effect was rescued by colonization with stable, defined minimal microbiota capable of SCFA production (OMM, n=6) (Fig. 5E), as well as liver regeneration, which was significantly reduced in GF mice (Fig. 5F). Even though the microbial composition of SPF controls is far more diverse compared to the OMM group (Fig. 5B, C, Suppl. Fig. 3), the general taxonomic distribution of SPF mice is well recapitulated and remains stable throughout liver regeneration in the OMM group, with major phyla being Firmicutes and Bacteroidota (Fig. 5D). Proliferation markers were assessed on mRNA level in liver tissue. Cyclins D1, A2 and B1, as well as the kinase CDK1, were hardly detectable in GF mice 48h after PHx in contrast to SPF mice, and at least partially rescued in OMM-colonized mice (Fig. 5G-H). No differences, however, were observed for cyclin E1. On protein level, phospho-retinoblastoma (RB) and CDK1 were strongly induced 48h after PHx in SPF mice, whereas GF mice lack detectable expression. OMM-mice show partially restored levels of CDK1 and phospho-RB (Fig. 5 I).

**Fig. 5.**
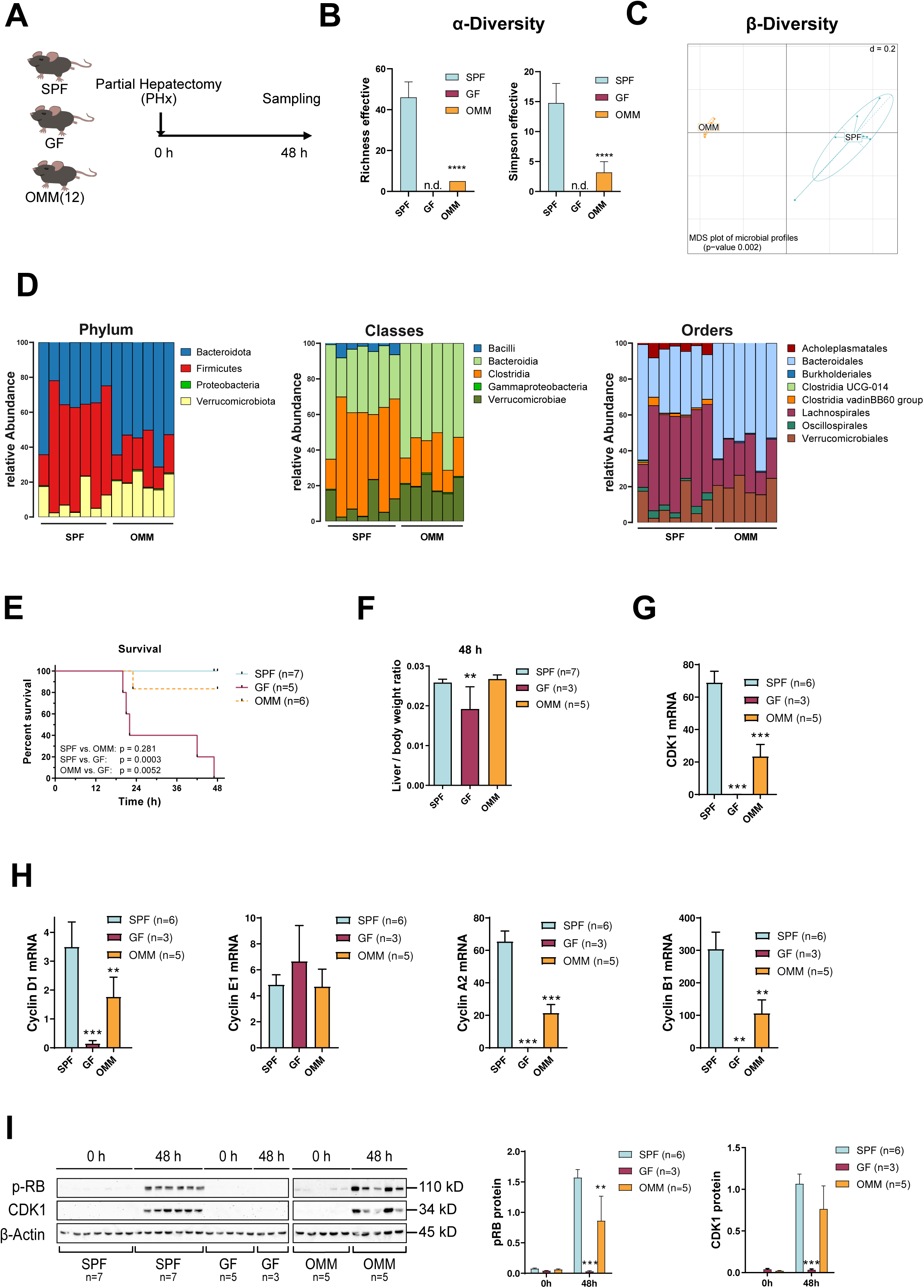
Liver regeneration, survival and hepatocyte proliferation after PHx are impaired in germfree mice, but rescued upon colonisation with a defined minimal bacterial consortium. (A) Partial hepatectomy (PHx) in control mice (specified pathogen free, SPF, n=7), germfree mice (GF, n=5), or gnotobiotic mice colonized with defined, stable minimal microbiota (OMM, n=6), and mice were sampled at the timepoint of PHx (0h), and after 48 h. (B) Diversity of microbial communities within the samples (α-diversity) based on V3-V4 16S rRNA sequencing are displayed as effective richness and Simpson effective. No values could be determined (n.d.) in GF mice. (C) Microbial diversity between SPF and OMM mice cecum content (β-diversity) is shown in a multidimensional scaling (MDS) plot. (D) Relative abundances of bacterial taxa at phylum, class, and order level display similar bacterial composition between SPF and OMM. (E) Kaplan-Meier survival analysis of SPF, GF and OMM mice after PHx. (F) Liver/body-weight ratio at 48h after PHx. Two mice in the GF group had to be euthanized before the 48h timepoint. (G, H) Cell cycle markers in liver tissue on mRNA (G) and protein level (H) show strongly impaired proliferation in GF liver parenchyma and partial rescue in OMM mice. ΔΔCt values normalized to β-actin housekeeping transcript, quantification of protein cell cycle markers phosphorylated RB and CDK1 at 0h and 48 h after PHx. .

### Hepatic lipid synthesis is blunted in germfree mice, but largely rescued by colonization with minimal bacterial consortium

As depicted schematically (Fig. 6A), hepatic lipid metabolism was traced by stable isotope labelling with two deuterated precursor compounds, [D9]-choline and [D4]- ethanolamine, administered by intraperitoneal injection. At 48h after PHx, GF mice showed essentially no incorporation of either [D9]-choline or [D4]-ethanolamine in de novo synthesized lipids. Controls (SPF) show robust lipogenic activity, coinciding with cell proliferation, whereas gnotobiotic OMM-mice show lipogenesis that is rescued at least partially, with largely conserved distribution of PC and PE species (Fig. 6B, C). As expected, the D4-label is preferentially incorporated in phospholipid species with long-chain polyunsaturated fatty acids (most prominent species: 38:6). Since mammalian liver can convert PE to PC by the phosphatidylethanolamine N-methyltransferase (PEMT), the amount of [D4]-label derived PC was quantified. The total amount of [D4]-labelled PC (Fig. 6D), was inferior to the [D9]-labelled fraction of PC (Fig. 6B) in the SPF group, in accordance with earlier findings. The most abundant [D4]-labelled PE species (38:6) was detected in similar amounts among the [D4]-labelled PC species, but not among the [D9]-labelled fraction of PC. This indicates a robust PE-PC conversion pathway in SPF controls which his essentially absent in GF mice, but rescued in the OMM-colonized group, with an equivalent amount of [D4]-labelled PC and similar lipid species profiles, compared to SPF mice. Analysis of labelled PE species from blood plasma confirmed these findings (Suppl. Fig. 3B, C). Next, expression of lipogenic enzymes in the liver (SCD1, FASN and ELOVL6) was analysed, and found to be downregulated at 48h after PHx, in accordance with experiments on antibiotics-treated mice (Fig. 6E). The desaturase SCD1 was also downregulated on protein levels at 48h, with heterogeneity (Fig. 6F, G). Compared to SPF controls, both GF and OMM-colonized mice had a significantly reduced expression of all three enzymes at the initial timepoint of hepatectomy. SCD1 and FASN showed higher relative expression in GF and OMM-mice at 48h compared to SPF controls, but at overall low levels (Fig. 6E).

**Fig. 6.**
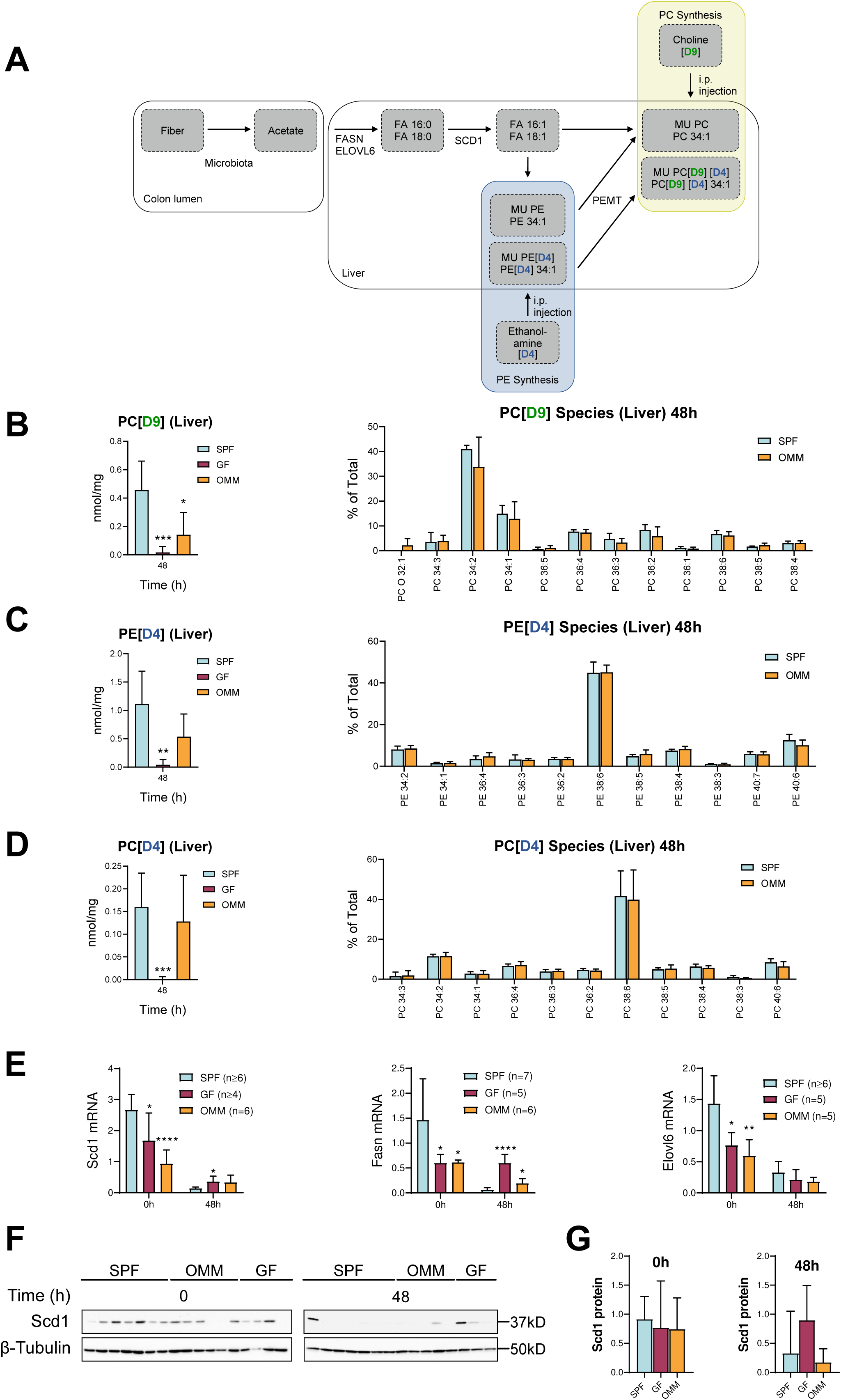
Phospholipid synthesis is impaired in germfree mice, but rescued in mice colonised with defined minimal bacterial consortium. (A) Schematic view: membrane lipid synthesis of stable isotope labeled D4 phosphatidyl ethanolamine (D4-PE), and D4 phosphatidyl choline (D4-PC), derived from labeled precursors D4-ethanolamine, and D9-choline. The hepatic enzyme phosphatidylethanolamine N-methyltransferase (PEMT) catalyzes the conversion from MU-PE to MU-PC. (B-D) Total amount of newly synthesized D9-PC (B), D4-PE (C) and D4-PC (D) measured by GC-MS/MS of liver tissue 2h after i.p. injection with D9- choline and D4-ethanolamine. Mean of lipid species distribution is normalized to the respective total amount of D9-PC, D4-PE or D4-PC in liver. (E) RNA levels of Fasn, Scd1 and Elovl5 were determined by qRT-PCR at 0h and 48h and normalized to β-actin. (F-G) Immunoblot of SCD1 and densitometric quantification.

### SCD1 expression is required for proliferation in human hepatomca cells and positively correlated with liver regeneration in patients

Since SCD1 was associated with gut microbiota, as well as with liver regeneration in mice, its functional contribution to liver cell proliferation was tested on human HepG2 hepatoma cells. SCD expression was downregulated by RNA interference with siRNA directed against all SCD isoforms, leading to significantly reduced expression of the cell cycle regulator cyclin A2 within 48h, compared to control siRNA (Fig. 7A, B). Accordingly, inhibition of SCD with a pharmacological compound significantly reduced cell proliferation, compared to mock-treated controls (Fig. 7C). Next, we assessed whether expression of lipogenic enzymes is associated with human liver regeneration. Three patients with liver metastasis of colorectal or cholangiocellular carcinoma, who underwent surgical intervention to increase the future liver remnant volume before tumour resection, were analysed. This intervention (associated liver partition and portal vein ligation for staged hepatectomy, or ALPPS) induces hypertrophy of non-diseased liver areas and hypotrophy of tumour-bearing areas. Radiological volumetry was carried out before ALPPS (pre intervention), after successful intervention but before tumour removal (post intervention), and after surgical tumour resection (post resection), shown for a representative patient (Fig. 7D). Biopsies from matched regenerating (hypertrophic, marked in red) and non-regenerating (hypotrophic, marked in blue) liver were obtained at the timepoint of tumour resection (Fig. 7D). SCD1 and FASN, but not ELOVL6, were increased on mRNA level in the hyperproliferative part of the liver in all three cases, as compared to the respective hypoproliferative tissue biopsy from the same patient (Fig. 7E). Due to limited group size, differences failed to attain significance. Next, expression levels in hyperproliferative, regenerating liver were correlated with the liver regeneration index, as determined by volumetry, either after intervention (Fig. 7F) or tumour resection (Fig. 7G). Expression of SCD1 and FASN, but not ELOVL6, was positively correlated with liver regeneration, attaining significance for SCD1 at the post-intervention timepoint (Fig. 7F).

**Fig. 7.**
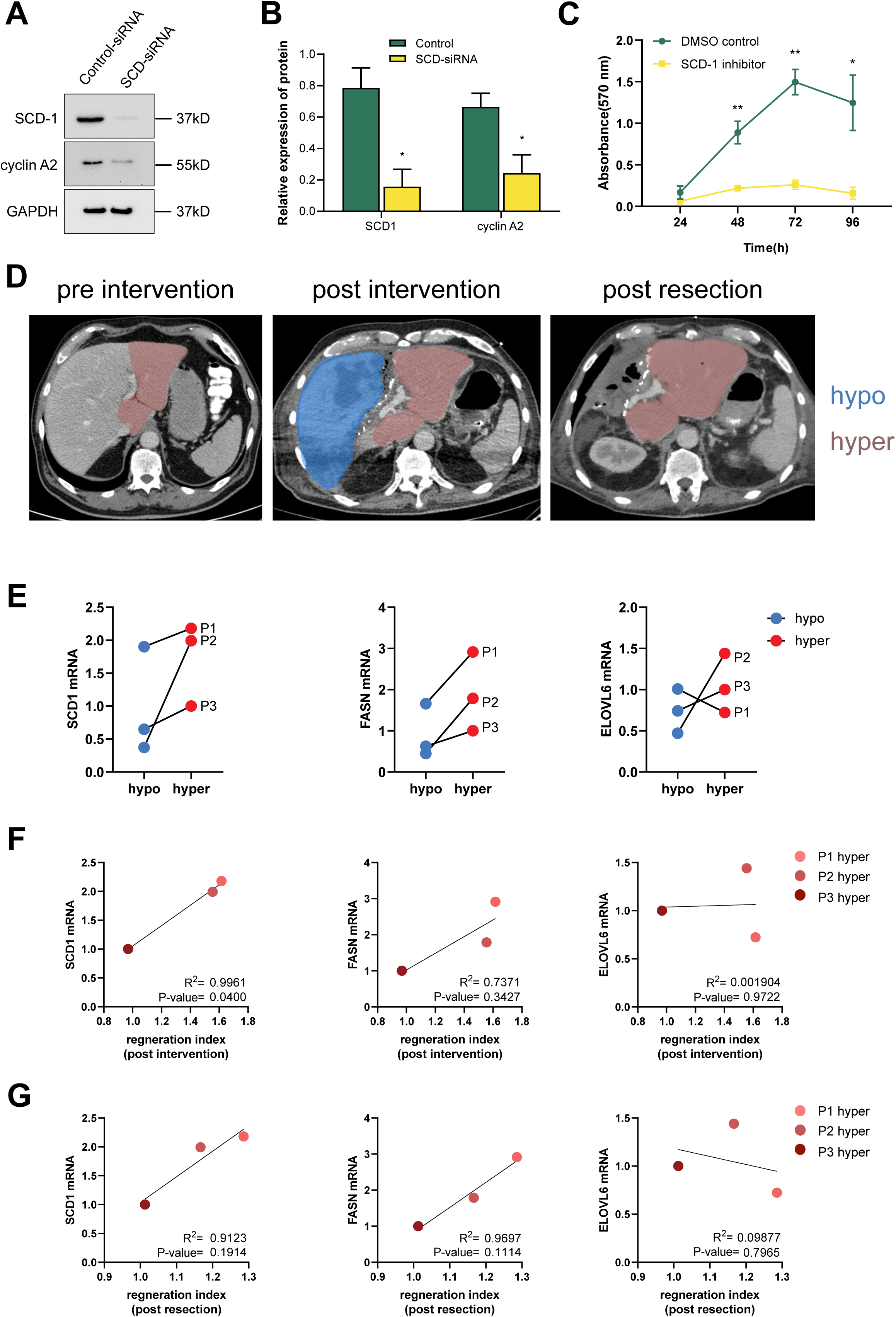
SCD1 contributes to proliferation in human hepatoma cells and is associated with liver proliferation and regeneration in human patients. (A and B) SCD1 and Cyclin A2 protein expression of HepG2 cell transfected with control or SCD-siRNA with GAPDH as a loading control with respective quantification. (C) Cell proliferation was measured by MTT assay at 24h, 48h, 72h and 96h for HepG2 cells treated with pharmacological SCD1 inhibitor, or mock-treated with DMSO. Mean absorbance of triplicate measurements. (D) CT scans of representative patient with hepatic metastasis of colon carcinoma undergoing PHx after pretreatment (ALPPS). Post intervention represents the timepoint immediately before surgical tumour resection. A biopsy of the remnant liver part (marked in red) was clinically described as hypertrophic and hyperplastic, the resected part (mareked in blue) as hypotrophic and hypoplastic. (E) Expression of key lipogenic enzymes in hypertrophic (red) compared to hypotrophic (blue) liver parenchyma biopsies in three individual patients, at the timepoint of resection. (F-G) SCD1 and FASN, but not ELOVL6 mRNA levels, measured in hyperproliferative tissue biopsies, are positively correlated with the volumetric regeneration index at timepoints post-intervention (F) or post tumour resection (G). Correlations were analysed with linear regression. Statistical significance was tested by two-tailed Student’s t test or Mann-Whitney U test. Data are presented as mean±SEM..

## Conclusions

Partial hepatectomy (PHx) is a major clinical procedure for patients with primary liver tumours, hepatic metastasis or parasitic cysts. Since the liver fulfills pivotal biochemical processes, it is necessary to quickly restore liver size and function after PHx to achieve optimal survival of patients. Consequently, it is essential to understand the mechanisms promoting or hindering liver regeneration, including extrahepatic factors delivered via the gut-liver axis. In fact, essential factors derived from gut microbiota affect liver regeneration,[34, 35] and gut microbiota have been associated with liver function and pathologies.[12, 14, 15, 36]

However, the functional link between microbial composition and liver regeneration, especially the effects of bacterial metabolites on hepatic lipid metabolism are still not fully understood. This system is potentially disturbed by gut dysbiosis, an obvious consequence of long-term antibiotics treatment, but also found in patients with liver cirrhosis or HCC.[36] Thus, we hypothesize that the detrimental effects of antibiotics- treatment on liver regeneration, which we report here in line with earlier findings,[37] may actually be attributable to alterations of hepatic lipid metabolism.

Within this study, a relatively short dysbiotic insult was induced applying broad-spectrum antibiotics in drinking water, which eliminate gram-negative and -positive bacteria.[38, 39] Since overgrowth by fungi is frequently observed [40, 41], the antimycotic fluconazole was added. Altogether, the treatment was well tolerated, in accordance with earlier reports[42]. The antibiotics treatment did not simply eradicate all bacteria but rather induced profound dysbiosis. In fact, significantly more aerobic colony forming units were found in in cecum content of antibiotics-treated mice. This may seem counterintuitive, yet it has been reported that use of antibiotics can increase the total number of bacteria while decreasing their diversity.[43–45] Dysbiosis produces a microaerophilic gut environment and promotes the growth of facultatively aerobic bacteria of the phylum Proteobacteria,[44] which usually make up only around 1% of the taxa.[10]

Thus, we hypothesize that specific shifts in the composition of gut bacteria, rather than globally decreased bacterial density, cause the observed massive problems for liver regeneration. Indeed, Proteobacteria could be observed in high abundance upon antibiotics treatment, whereas obligate anaerobic bacteria of the phyla Bacteroidetes and Firmicutes were strongly diminished. This is a crucial finding regarding microbial functionality, since Firmicutes and Bacteroidetes typically ferment dietary fiber to short chain fatty acids, in contrast to Proteobacteria.[10, 35, 44] Thus, absence of fermentation-competent bacteria may lead to a lack of microbial-derived metabolites that are crucially required for hepatic lipogenesis and liver cell growth and proliferation. Interestingly, some antibiotics-treated mice displayed an increase in relative abundance of the fermenting genera Clostridiales and Lachnospirales at late timepoints after PHx, in line with a concomitant increase in hepatocyte proliferation. This effect may correspond to adaptation of the microbiota after longer treatment.[41]

Whereas several studies monitored the microbial shifts associated with liver regeneration after partial hepatectomy in rodent models, a clear picture failed to emerge, likely due to differences in experimental design.[46, 47] In the present study, microbiota shifted to a higher relative abundance of Firmicutes and a decrease in Bacteroidetes at 48h after PHx, but returned to initial levels after 96h. In the antibiotics-treated group, a slight but significant decrease in liver weight was observed at the timepoint of hepatectomy, in accordance with earlier findings.[11] However, no signs for liver toxicity or putative damage caused by antibiotics-treatment were observable histologically, in line with published findings showing that the treatment is generally well-tolerated, even for longer periods of several weeks.[20, 38]

Importantly, liver regeneration was severely delayed in the antibiotics-treated group, and survival of mice after PHx was compromised. In accordance, there was a drastic lag in hepatocyte proliferation: expression of most of the crucial cell cycle markers was only detected one week after PHx in the dysbiotic mice, confirmed by histological Ki67 staining. Of note, the data did not suggest an arrest in a specific cell cycle phase, but rather a general delay of proliferation in the antibiotics-treated group. This delayed proliferation peak appeared at the same timepoint when fermentation-competent bacteria were again detectable in some of the antibiotics-treated mice.

Thus, it can be assumed that the severely impaired liver regeneration and hepatocyte proliferation in antibiotics-treated mice are due to the imbalance of gut microbiota. We hypothesize that the transition from quiescence to cell cycle entry may have been delayed in hepatocytes, due to lack of general metabolic factors, such as lipids. Hepatocytes in antibiotic treated mice were actually found to be smaller in size compared to control mice upon histological analysis, supporting the assumption that cell growth and proliferation are blunted, already during the early priming phase of regeneration. Indeed, proliferation and lipid synthesis have been shown to be closely linked in other model organisms.[48–51]

Thus, we focused here on short-chain fatty acids (SCFA), as a putative link between gut dysbiosis and impaired hepatocyte proliferation. SCFA are derived from intestinal microbial fermentation of dietary fiber,[10] and are primarily absorbed through the portal vein during lipid digestion.[52] SCFA play a variety of physiological roles: butyrate is the primary energy source for colonocytes, while the liver mainly requires acetate.[53, 54] Furthermore, hepatic lipogenesis, which produces de novo lipids that are crucial not only for hepatocyte growth and proliferation, but also for systemic lipid metabolism, relies on bacterial-derived SCFA. Here, we could show that the concentration of all SCFA species was significantly reduced in gut content after antibiotics treatment, and throughout the duration of the experiment. Acetate, as the most abundant SCFA, is used as building block for saturated fatty acid (FA) synthesis by the enzyme FASN.[11] The saturated fatty acids are further modified in hepatocytes by the desaturase SCD1 to produce monounsaturated fatty acids (MUFA, e.g. palmitoleic or oleic acid), and/or processed by elongase enzymes, and hence incorporated in cellular membranes as phospholipids. In accordance, stable isotope labelling experiments, focussing on monounsaturated phosphatidylcholine (MU-PC) as most important membrane lipid component, clearly showed that the de novo synthesis of phospholipids was significantly decreased in the liver of antibiotics treated mice after hepatectomy, even more clearly observable in blood plasma.

This model, linking bacterially derived SCFA with liver lipid metabolism and regeneration, was tested next. A significant association between acetate levels in the gut and liver regeneration (as liver-to-body-weight ratio) was confirmed. Further, acetate levels in the gut were highly significantly associated with host lipogenesis, evidenced by quantification of labeled MU-PC in the blood. Lastly, systemic lipogenesis, evidenced as labelled MU-PC in the blood plasma, was also significantly associated with liver regeneration. Furthermore, the expression of host enzymes involved in fatty acid synthesis and modification was tested in liver tissue. The desaturase SCD1, but not FASN or ELVOL6, showed a clear dependence on the gut microbiome status, in accordance with our earlier findings,[11] and its expression kinetics on RNA and protein levels coincided with proliferation markers after PHx.

However, the data on antibiotics treated mice do not allow to establish causality and may be prone to inherent heterogeneity of the dysbiotic insults. Therefore, we have tested whether a minimal set of microbiota, capable of SCFA production, is sufficient for efficient liver regeneration. First, the partial hepatectomy model was established on gnotobiotic germfree mice, by focusing on the timepoint of 48h. Germfree mice (GF) showed severely impaired liver regeneration and high mortality after PHx. Similar to the antibiotics treated group at early timepoints, cell proliferation was severely blocked. Importantly, all of these effects were essentially rescued upon colonization of GF mice with a stable and defined minimal set of microbial taxa that is capable of SCFA production.[11, 19]

Thus, hepatocyte proliferation, liver regeneration and associated mortality were fully restored by re-introducing a defined minimal microbiome that resembles the conventionally maintained controls (SPF) on phylum, class and order level. Of course, it cannot be absolutely excluded that bacterial contaminations may occur during the PHx intervention, despite sterile conditions during surgery. However, 16S rRNA gene analysis confirmed that GF and OMM mice retained their respective gut microbiota status at the 48h timepoint after PHx, with no bacterial taxa detectable in any GF mouse. Next, we have assessed the putative consequences on lipid metabolism. GF mice showed largely blunted de novo lipogenesis in the liver, as evidenced by tracing stable isotope labelled phosphatidylethanolamine and phosphatidylcholine levels, in good accordance with earlier findings.[11] Phospholipid synthesis was at least partially restored in the OMM group, coinciding with the restoration of liver regeneration and hepatocyte proliferation. The expression of lipogenic enzymes was found to be more heterogeneous, but generally decreased in GF and OMM groups. SCD1 levels were decreased at 48h after PHx, similar to the observations after antibiotics treatment.

Based on the finding that the expression of SCD1 was differentially regulated during liver regeneration in mice, we tested whether its expression is required for cell proliferation in human hepatoma cells. Pharmacological SCD1 inhibition or downregulation by RNA-interference indeed lead to significantly reduced cell proliferation in vitro.

Finally, we assessed whether the findings from animal models have any translational relevance, especially regarding the putative association of lipid metabolism with liver regeneration. As surrogate markers, we assessed the expression lipogenic enzymes in human liver tissue biopsies, taking advantage of matched parallel liver biopsies from individual patients undergoing a specific surgical intervention to increase the future liver remnant (FLR). In fact, patients with primary liver cancer or hepatic metastases often require extensive hepatic resection due to the large size of the tumour(s) adjacent to important vascular structures, resulting in an insufficient volume of the remaining liver. The generally accepted safe minimum threshold for the FLR varies from 20 to 25% for a normal liver, to >40% for a cirrhotic liver.[55–57] Thus, ensuring adequate FLR is one of the key factors in determining liver tumour resection, and methods currently used to increase FLR volume include associated liver partition and portal vein ligation for staged hepatectomy (ALPPS). We thus selected three oncological patients who received ALPPS treatment to achieve sufficient FLR before liver resection. This preoperative surgical intervention induces hyperproliferative as well as atrophic regions within an individual liver, and biopsies from these tissue areas were obtained for analysis. Expression of SCD1 and FASN, but not the elongase ELOVL6, was increased in hyperproliferative areas in all three cases, as compared to hypoproliferative parts from the same patient. Further, radiological assessment of liver regeneration by volumetry showed a positive correlation between expression of SCD1 and FASN with the recovery level of liver regeneration at two critical timepoints: after completion of the ALPPS intervention, and after the tumour resection. Thus, expression of lipogenic enzymes is positively associated with liver proliferation and regeneration in humans.

From a translational perspective, the findings on preclinical models presented here may have important repercussions for human patients who depend on efficient liver regeneration, e.g., after oncological resections. Dysbiosis could arise frequently in patients due to antibiotics treatment or liver cirrhosis, persistently disturbing the gut microbiota for several months. As a consequence, the dysbiotic gut microbiota may fail to produce sufficient amounts of SCFA, required as pivotal building blocks for membrane lipid synthesis in the liver. Hence, liver regeneration and even patient survival could be directly linked to the composition of gut microbiota. These findings may offer new avenues for preoperative screening by analyzing microbial composition via well established 16S rRNA gene sequencing, or even therapeutic interventions. However, interfering with gut microbiota is certainly not straightforward. Of note, whereas SCFA are beneficial for liver function, they were also connected to HCC formation, albeit at high concentrations.[58] Therefore, more studies are needed to fully understand the connection between bacterial fermentation, lipid synthesis and liver regeneration.

## Abbreviations

ABX: antibiotics-treated
Cer: Ceramide
FA: Fatty acid
FASN: Fatty acid synthase gene
GF: Germfree
i.p.: Intraperitoneal injection
LPC: Lyso-phosphatidylcholine
LPE: Lyso-phosphatidylethanolamine
MAG: Monoacylglycerol
MU PC: Phosphatidylcholine with monounsaturated acyl side chain
OMM: Mice stably colonised with twelve major intestinal bacteria taxa, housed in a gnotobiotic environment
PC: Phosphatidylcholine
PE: Phosphatidylethanolamine
PEMT: Phosphatidylethanolamine N-methyltransferase
PE P: PE-based plasmalogens
PG: Phosphatidylglycerol
PHX: partial hepatectomy
PI: Phosphatdiylinositol
PS: Phosphatidylserine
SCD1: Steaoryl-Co-A-Desaturase
SPF: Specific pathogen free
TG: Triacylglycerol

## Acknowledgements

The authors wish to thank

## Authors’ contributions

Study conception and design (BH, JE, NH, KPJ), performed experiments (YY, AS, XJZ, JE, ML, GL, MH, FL, YG), provided materials (BH, MB, AB, HF), data analysis and interpretation (AS, JE, ML, GL, MH, FL, HF, CM, DH, BH, NH, KPJ), wrote manuscript (YY, AS, KPJ), obtained funding (JE, KPJ).

---

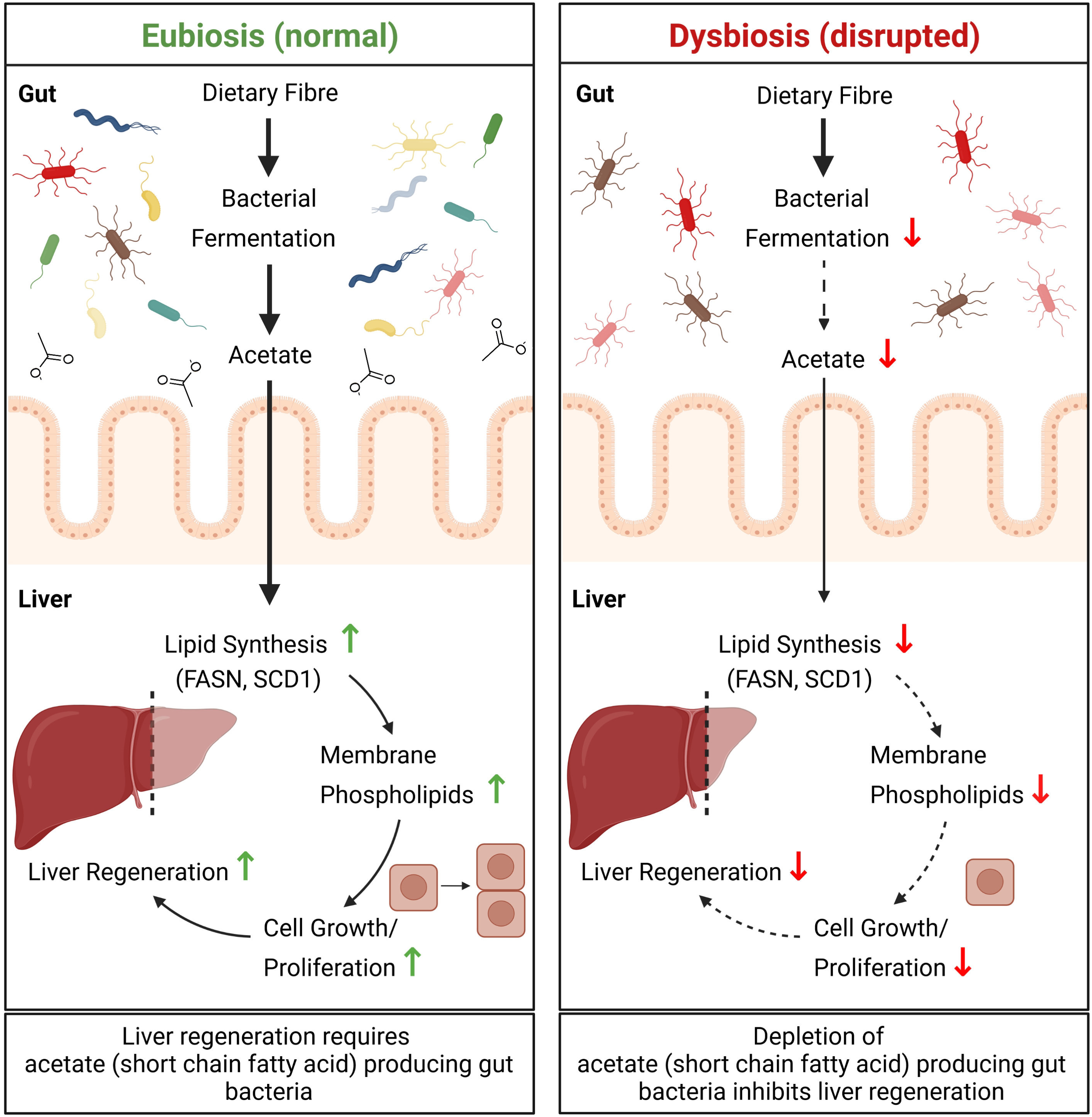

