## Supplementary Material for "The gut microbiota promotes liver regeneration through hepatic membrane phospholipid synthesis"

**Table of contents:**

Supplementary Methods

Supplementary Figure Legends

Supplemtentary Figures

Supplementary Tables

The Supplementary material should have a manuscript title, list of authors, a table of contents, followed by the list of investigators (if there is one), text (such as methods), figures, tables, and then references. Supplementary material must be prepared as a single Word file with pages numbered (including references, tables and figure legends) using Times New Roman or Arial 12 pt double-spaced. Sections have to be 12 pt bold, subsections have to be 12 pt, italics.

**Supplementary Figure Legends**

**Suppl. Fig. 1.** (A) Serial dilutions aerobically grown on standard blood agar plates, pictures take after over night culture at 37 °C of feces (left) collected during PHx or cecal content (right) collected after sampling of mice (B) Liver/body weight ratio is decreased in antibiotics-treated mice after three days. (C) Liver/body weight ratios were normalized to the respective values at 0h timepoint, and calculated accordingly for all following time points after PHx (n=6-10 mice per time point and group). (D) Representative immunohistochemical staining images of Ki67-positive hepatocytes of tissue sections from antibiotics-treated (ABX) or control (Ctrl) mice. Scale bars represent 50 µm. (E) Visual quantification of hepatocyte numbers per $\mu m^2$ on H&E stained tissue sections in control and antibiotics-treated mice (n=6 mice per time point and group), mean ± SEM. Higher number of hepatocytes per area indicate smaller cell size.

**Suppl. Fig. 2.** Quantity of propionate (A), butyrate (B), iso-butyrate (C), and total short chain fatty acids (SCFA)(D) in colon content of antibiotics-treated (red) versus control mice (blue). Analysis by GC-MS/MS, shown in all cases mean ± SD. Correlation of colonic acetate levels with liver/bodyweight of mice at different timepoints after PHx (E), and correlation of colonic acetate levels with cyclin E1 RNA expression in liver parenchyma (F). (G-I) Hepatic stable isotope labeled phosphatidylcholine (PC[D9]) species, containing one monounsaturated acyl chain, 32:1 (G), 34:1(H), and 36:1 (I). (J-L) Monounsaturated PC[D9] species measured in plasma of ABX (red) and control mice (blue). (M-O) Correlation of de novo synthesized monounsaturated D9 stable isotope labeled phosphatidyl choline (MU-PC[D9]) in liver with colonic acetate (M), with liver/bodyweight ratio (N), and normalized liver/bodyweight ratio (O) of the timepoints 0, 24, 48 and 72 h. (P) Plasmatic MU-PC[D9] levels correlated with liver/bodyweight ratio. (Q-R) Correlation of total MU-PC[D9] in liver (Q) or blood plasma (R) with mRNA levels of cyclin E1 in hepatic parenchyma. For correlation, linear regression was applied. Statistical analysis was conducted using unpaired two-tailed student's t-test, with *P < 0.05, **P <0.01, and ***P < 0.001.

**Suppl. Fig. 3.** (A) Gel electrophoretic analysis of PCR fragments of V3-V4 variable regions of 16S genomic DNA, isolated from cecal content of specific pathogen free (SPF), Oligo Mouse Microbiota 12 (OMM) colonized and germ free mice (GF), with 1 kb DNA Ladder (M), negative control without added template DNA (- ctrl) and positive control (+ ctrl). (B) As expected, the relative abundances of bacterial taxa differ at the taxonomic genus level between SPF controls and OMM-colonized mice, based on 16S sequencing data. (C) In germfree mice, incorporation of stable isotope labelled phosphatidylethanolamine [D4-PE] after PHx is blunted, but essentially rescued in OMM-colonized mice, as shown by detection of total labelled PE lipids in blood plasma (D). The lipid species distribution for PE species detected in blood plasma at 48h after PHx does not differ between SPF control mice and OMM-controls (right panel). Essentially the same is observed for detection of stable isotope labelled phosphatidylcholine [D9-PC] after PHx, which is almost absent in germfree mice, but essentially rescued in OMM-colonized mice (D), with a highly conserved lipid species profile in OMM-mice, compared to controls (right panel).

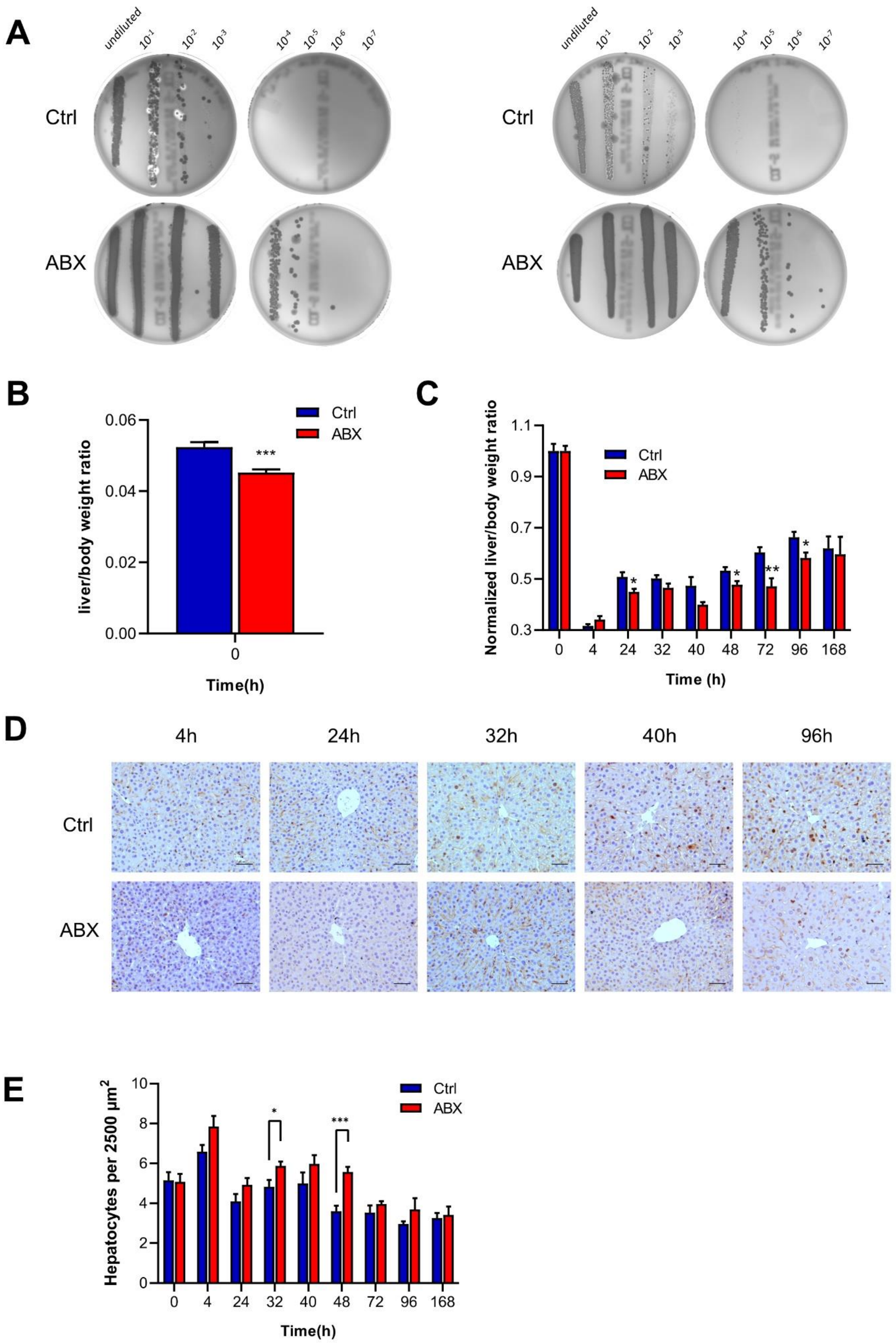
A
undiluted
$10^{-1}$
$10^{-2}$
$10^{-3}$
$10^{-4}$
$10^{-5}$
$10^{-6}$
$10^{-7}$
Ctrl
ABX
B
liver/body weight ratio
Ctrl
ABX
***
Time(h)
C
Normalized liver/body weight ratio
Time (h)
D
4h
24h
32h
40h
96h
E
Hepatocytes per 2500 μm²
Time(h)

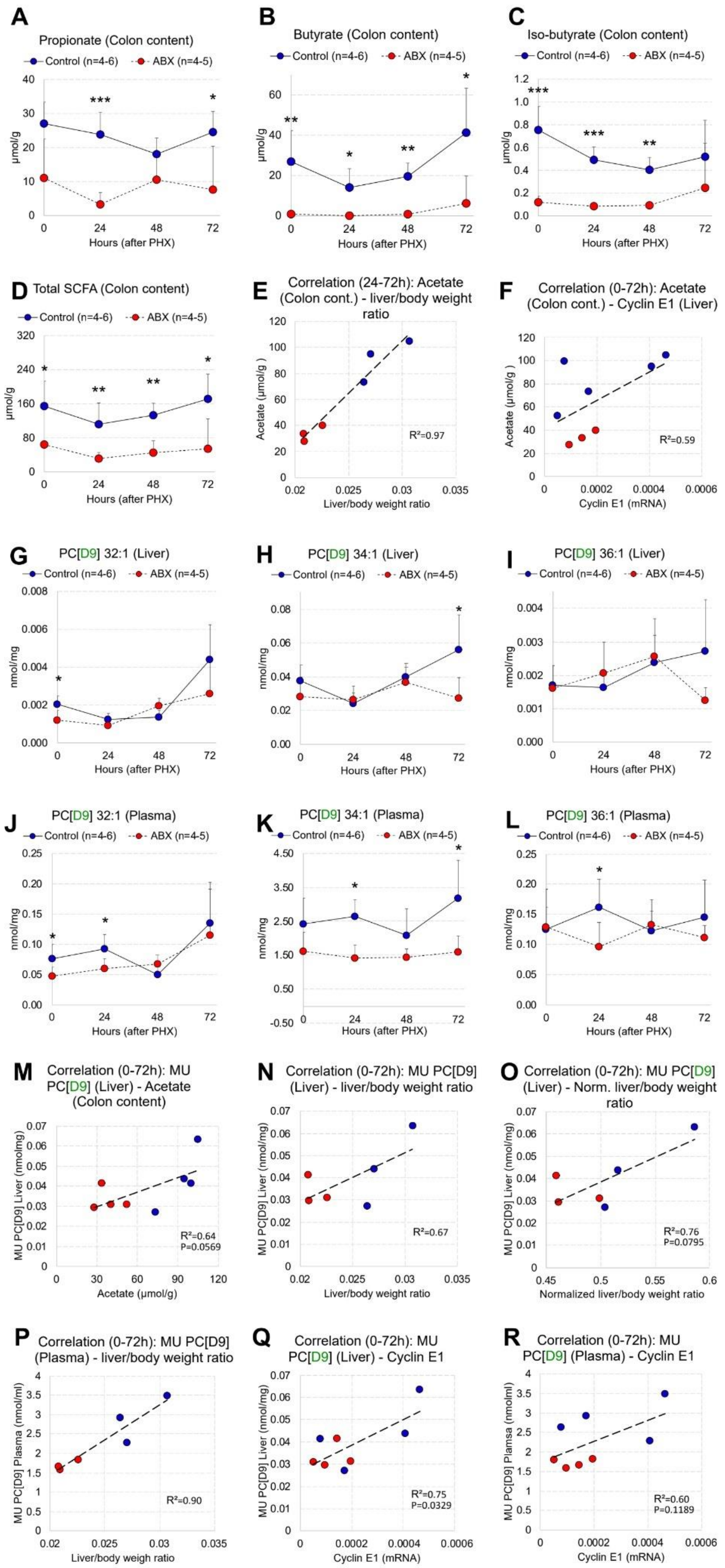
A
Propionate (Colon content)
Control (n=4-6)
ABX (n=4-5)
µmol/g
Hours (after PHX)
B
Butyrate (Colon content)
C
Iso-butyrate (Colon content)
D
Total SCFA (Colon content)
E
Correlation (24-72h): Acetate (Colon cont.) - liver/body weight ratio
Acetate (µmol/g)
Liver/body weight ratio
R²=0.97
F
Correlation (0-72h): Acetate (Colon cont.) - Cyclin E1 (Liver)
Cyclin E1 (mRNA)
R²=0.59
G
PC[D9] 32:1 (Liver)
nmol/mg
H
PC[D9] 34:1 (Liver)
I
PC[D9] 36:1 (Liver)
J
PC[D9] 32:1 (Plasma)
K
PC[D9] 34:1 (Plasma)
L
PC[D9] 36:1 (Plasma)
M
Correlation (0-72h): MU PC[D9] (Liver) - Acetate (Colon content)
MU PC[D9] Liver (nmol/mg)
Acetate (µmol/g)
R²=0.64
P=0.0569
N
Correlation (0-72h): MU PC[D9] (Liver) - liver/body weight ratio
Liver/body weight ratio
R²=0.67
O
Correlation (0-72h): MU PC[D9] (Liver) - Norm. liver/body weight ratio
Normalized liver/body weight ratio
R²=0.76
P=0.0795
P
Correlation (0-72h): MU PC[D9] (Plasma) - liver/body weight ratio
MU PC[D9] Plasma (nmol/ml)
Liver/body weigh ratio
R²=0.90
Q
Correlation (0-72h): MU PC[D9] (Liver) - Cyclin E1
Cyclin E1 (mRNA)
R²=0.75
P=0.0329
R
Correlation (0-72h): MU PC[D9] (Plasma) - Cyclin E1
MU PC[D9] Plamsa (nmolml)
R²=0.60
P=0.1189

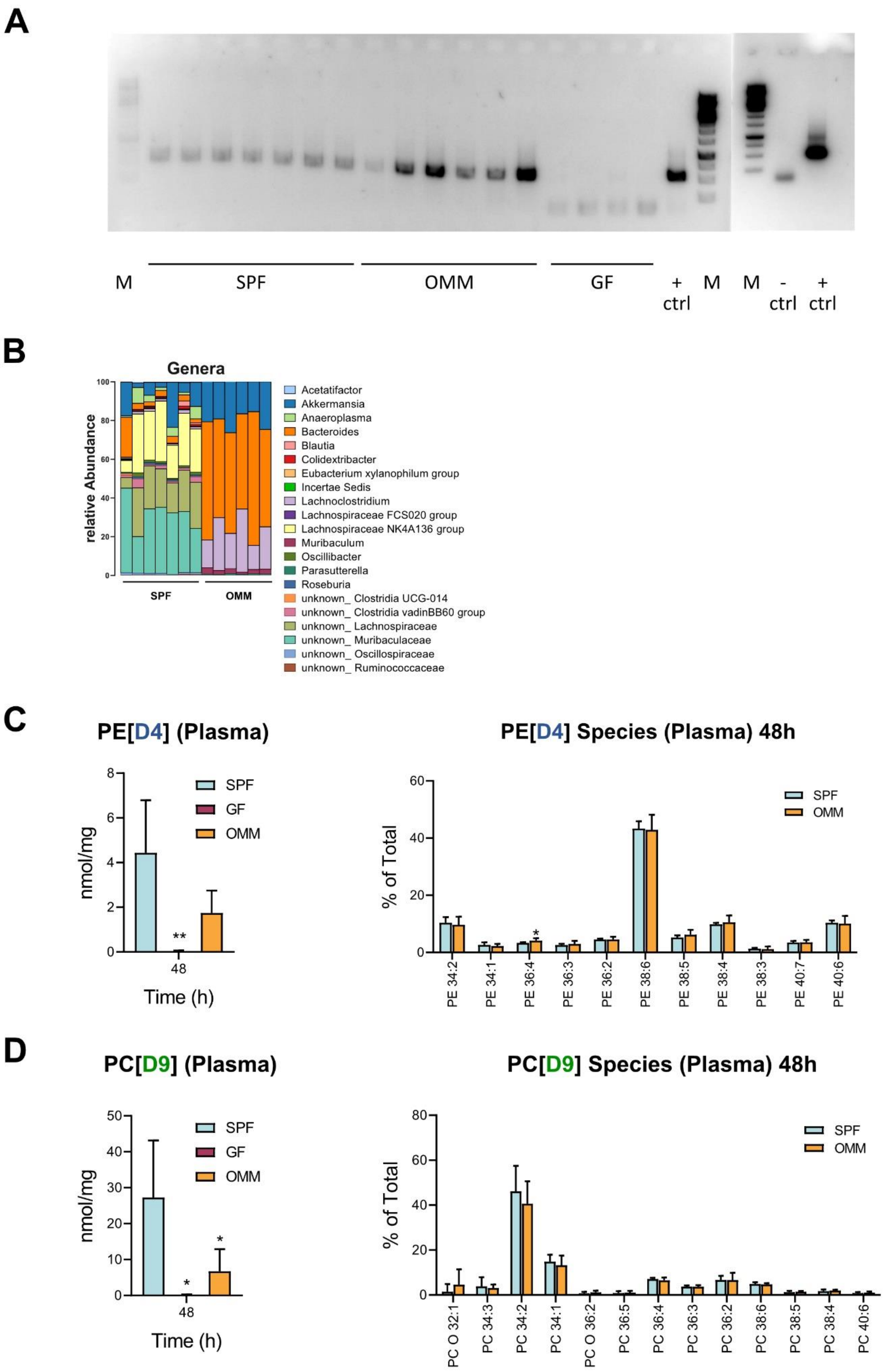
A
M
SPF
OMM
GF
+ ctrl
M
M
- ctrl
+ ctrl
B
Genera
relative Abundance
SPF
OMM
Acetatifactor
Akkermansia
Anaeroplasma
Bacteroides
Blautia
Colidextribacter
Eubacterium xylanophilum group
Incertae Sedis
Lachnoclostridium
Lachnospiraceae FCS020 group
Lachnospiraceae NK4A136 group
Muribaculum
Oscillibacter
Parasutterella
Roseburia
unknown_ Clostridia UCG-014
unknown_ Clostridia vadinBB60 group
unknown_ Lachnospiraceae
unknown_ Muribaculaceae
unknown_ Oscillospiraceae
unknown_ Ruminococcaceae
C
PE[D4] (Plasma)
nmol/mg
Time (h)
SPF
GF
OMM
PE[D4] Species (Plasma) 48h
% of Total
PE 34:2
PE 34:1
PE 36:4
PE 36:3
PE 36:2
PE 38:6
PE 38:5
PE 38:4
PE 38:3
PE 40:7
PE 40:6
D
PC[D9] (Plasma)
nmol/mg
Time (h)
PC[D9] Species (Plasma) 48h
% of Total
PC O 32:1
PC 34:3
PC 34:2
PC 34:1
PC O 36:2
PC 36:5
PC 36:4
PC 36:3
PC 36:2
PC 38:6
PC 38:5
PC 38:4
PC 40:6

**Supplementary Table 1:**
List of Antibodies used

| **Target Epitope** | **Species reactivity** | **Reference Number** | **Provider** |
|---|---|---|---|
| cyclin B1 | Murine | #4138S | Cell Signaling |
| cyclin A2 | Murine | ab181591 | Abcam |
| phosphorylated retinoblastoma protein (p-RB)(Ser807/811) | Murine | #8516S | Cell Signaling |
| CDK1 | Murine | ab131450 | Abcam |
| Ki67 | Murine | | BD Pharmingen |
| SCD1 | Human | ab19862 | Abcam |
| SCD1 | Murine | #2794 | Cell Signaling |
| β-actin | Murine | #3700 | Cell Signaling |
| β-tubulin | Murine | ab6046 | Abcam |
| GAPDH | Murine | sc32233 | Santa Cruz |
| GAPDH | Human | #2118 | Cell Signaling |

**Supplementary Table 2**: Oligonucleotide Primers

| Target | Sequences | Species | Application |
|---|---|---|---|
| Ccna2 (cyclin A2) sense antisense | 5'-CTT GGC TGC ACC AAC AGT AA-3' 5'-CAA ACT CAG TTC TCCCAA AAA CA-3' | murine | RT-qPCR KAPA SYBR FAST Kit |
| CcnB1 (cyclin B1) sense antisense | 5'-GCT TAGCGC TGA AAA TTC TTG-3' 5'-TCT TAG CCA GGT GCT GCA TA-3' | murine | RT-qPCR KAPA SYBR FAST Kit |
| Ccnd1 (cyclin D1) sense antisense | 5'-TTT CTT TCC AGA GTC ATC AAG TGT-3' 5'-TGA CTC CAG AAG GGCTTC AA-3' | murine | RT-qPCR KAPA SYBR FAST Kit |
| Ccne1 (cyclin E1) sense antisense | 5'-TTT CTG CAG CGT CAT CCT C-3' 5'-TGG AGC TTA TAG ACT TCG CAC A-3' | murine | RT-qPCR KAPA SYBR FAST Kit |
| Scd1 sense antisense | 5'-TTC CCT CCT GCA AGC TCT AC-3' 5'-CAG AGC GCT GGT CAT GTA GT-3' | murine | RT-qPCR Universal Probe Library (UPL) Probe #34 |
| Fasn sense antisense | 5'-GCT GCT GTT GGA AGT CAG C-3' 5'-AGT GTT CGT TCC TCG GAG TG-3' | murine | RT-qPCR UPL Probe #58 |
| Elovl6 sense antisense | 5'-CAG CAA AGC ACC CGA ACT A-3' 5'-AGG AGC ACA GTG ATG TGG TG-3' | murine | RT-qPCR UPL Probe #4 |
| Actb sense antisense | 5'-AAG GCC AAC CGT GAA AAG AT-3' 5'- GTG GTA CGA CCA GAG GCA TA-3' | murine | qPCR KAPA SYBR FAST Kit UPL Probe #22 |
| Gapdh sense antisense | 5'-AGG TGG GTG TGA ACG GAT TTG-3' 5'-TGT AGA CCA TGT AGT TGA GGT CA-3' | murine | qPCR KAPA SYBR FAST Kit |
| HPRT sense antisense | 5’-GAC CAG TCA ACA GGG GAC AT-3’ 5’-GTG TCA ATT ATA TCT TCC ACA ATC AAG-3’ | human | qPCR UPL Probe #22 |
| SCD1 sense antisense | 5’-CCT ACC TGC AAG TTC TAC ACC TG-3’ 5’-GAC GAT GAG CTC CTG CTG TT -3’ | human | qPCR UPL Probe #37 |
| ELOVL6 sense antisense | 5’-CAA AGC ACC CGA ACT AGG AG -3’ 5’-GGT GAT ACC AGT GCA GGA AGA-3’ | human | qPCR UPL Probe #38 |
| FASN sense antisense | 5’-CAG GCA CAC ACG ATG GAC-3’ 5’-CGG AGT GAA TCT GGG TTG AT-3’ | human | qPCR UPL Probe #11 |
